# Activated α_IIb_β_3_ on platelets mediates flow-dependent NETosis via SLC44A2

**DOI:** 10.1101/373670

**Authors:** Adela Constantinescu-Bercu, Luigi Grassi, Mattia Frontini, Isabelle I. Salles-Crawley, Kevin J Woollard, James T.B. Crawley

## Abstract

Platelet-neutrophil interactions are important for innate immunity, but also contribute to the pathogenesis of deep vein thrombosis, myocardial infarction and stroke. Here we report that, under flow, von Willebrand factor/glycoprotein Ibα-dependent platelet ‘priming’ induces integrin α_IIb_β_3_ activation that, in turn, mediates neutrophil and T-cell binding. Binding of platelet α_IIb_β_3_ to SLC44A2 on neutrophils leads to mechanosensitive-dependent production of highly prothrombotic neutrophil extracellular traps. A polymorphism in *SLC44A2* (rs2288904-A) present in 22% of the population causes an R154Q substitution in an extracellular loop of SLC44A2 that is protective against venous thrombosis results in severely impaired binding to both activated α_IIb_β_3_ and VWF-primed platelets. This was confirmed using neutrophils homozygous for the *SLC44A2* R154Q polymorphism. Taken together, these data reveal a previously unreported mode of platelet-neutrophil cross-talk, mechanosensitive NET production, and provide mechanistic insight into the protective effect of the *SLC44A2* rs2288904-A polymorphism in venous thrombosis.

**Summary:** Platelets that are primed following interaction with von Willebrand factor under flow mediated direct interactions with neutrophils via activated platelet integrin, α_IIb_β_3_, and SLC44A2 on neutrophils. This interaction initiates signaling in a mechanosensitive manner that promotes neutrophil extracellular trap formation.

## Introduction

To fulfil their hemostatic function, platelets must be recruited to sites of vessel damage. This process is highly dependent upon von Willebrand factor (VWF). Upon vessel injury, exposed subendothelial collagen binds plasma VWF via its A3 domain (Cruz et al., 1995). Elevated shear, or turbulent/disturbed flow, then unravels tethered VWF and exposes its A1 domain, facilitating specific capture of platelets via glycoprotein (GP)Ibα.

As well as capturing platelets under flow, the A1-GPIbα interaction also induces shear-dependent signaling events (Bryckaert et al., 2015). For this, GPIbα first binds the A1 domain of immobilized VWF (Zhang et al., 2015). Rheological forces then cause unfolding of the GPIbα mechanosensitive domain that translates the mechanical stimulus into a signal within the platelet (Ju et al., 2016; Zhang et al., 2015). This leads to release of intracellular Ca^2+^ stores and activation of the platelet integrin, α_IIb_β_3_ (Gardiner et al., 2010).

VWF-mediated signaling transduces a mild signal. Consequently, these signaling events are often considered redundant within hemostasis as platelets respond more dramatically to other agonists present at sites of vessel injury (e.g. collagen, thrombin, ADP, thromboxane A2) (Jackson et al., 2003; Senis et al., 2014). Full platelet activation involves release of α- and δ-granules, presentation of new cell surface proteins, activation of cell surface integrins and alterations in the membrane phospholipid composition. The extent of platelet activation is dependent upon both the concentration, and identity, of the agonist(s) to which the platelets are exposed, which is dictated by the location of the platelets relative to the damaged vessel. For example, platelets in the core of a hemostatic plug/thrombus are exposed to higher concentrations of agonists and are more highly activated (i.e. P-selectin-positive procoagulant platelets) than those in the surrounding shell (P-selectin-negative) (de Witt et al., 2014; Shen et al., 2017; Stalker et al., 2013; Welsh et al., 2014). Thus, platelets exhibit a ‘tunable’ activation response determined by agonist availability.

Aside from hemostasis, platelets also have important roles as immune cells by aiding in targeting of bacteria by leukocytes (Gaertner et al., 2017; Kolaczkowska et al., 2015; Sreeramkumar et al., 2014; Wong et al., 2013). Platelet-leukocyte interactions also influence the development of inflammatory cardiovascular conditions. In deep vein thrombosis (DVT), VWF-dependent platelet recruitment, platelet-neutrophil interactions and the production of highly thrombotic neutrophil extracellular traps (NETs) all contribute to the development of a pathological thrombus (Brill et al., 2011; Brill et al., 2012; Fuchs et al., 2012a; Schulz et al., 2013; von Bruhl et al., 2012). Although the precise sequence of events still remains unclear, it appears that during the early stages of DVT, VWF-bound platelets acquire the ability to interact with leukocytes (von Bruhl et al., 2012). Exactly how this is mediated given the lack of vessel damage is unclear. It also remains to be determined precisely how platelet-tethered neutrophils undergo NETosis in DVT in the absence of an infectious agent.

Known direct platelet-leukocyte interactions involve either P-selectin or CD40L on the surface of platelets binding to P-selectin glycoligand-1 (PSGL-1) and CD40, respectively, on leukocytes (Lievens et al., 2010; Mayadas et al., 1993; Palabrica et al., 1992). As platelets must be potently activated to facilitate P-selectin/CD40L exposure, such interactions unlikely mediate the early platelet-leukocyte interactions that occur in the murine DVT model. Consistent with this, lack of platelet P-selectin has no effect upon either leukocyte recruitment or thrombus formation in murine DVT (von Bruhl et al., 2012). Leukocytes can also indirectly interact with platelets through Mac-1 (integrin α_M_β_2_), which can associate with activated α_IIb_β_3_ via fibrinogen (Weber and Springer, 1997), or directly via GPIbα (Simon et al., 2000). Interactions are also possible through lymphocyte function-associated antigen 1 (LFA-1/integrin α_L_β_2_) that can bind intercellular adhesion molecule 2 (ICAM-2) on platelets (Damle et al., 1992; Diacovo et al., 1994). In both instances though, leukocyte activation is necessary to activate Mac-1 or LFA-1 integrins before interactions can occur.

Although it is often assumed that only activated platelets bind leukocytes, recent studies have revealed that platelets captured under flow by VWF released from activated endothelial cells can recruit leukocytes (Doddapattar et al., 2018; Zheng et al., 2015). If VWF-GPIbα-dependent signaling is capable of promoting leukocyte binding, this may be highly relevant to the non-hemostatic platelet functions (particularly when other agonists are not available/abundant), but may also provide major mechanistic insights into the early recruitment of leukocytes during the initiation of DVT.

Genome wide association studies (GWAS) on venous thromboembolism (VTE) have identified a panel of genes (*ABO, F2, F5, F11, FGG, PROCR*) with well-described influences upon coagulation and thrombotic risk, as well as those with well-established causative links (e.g. *PROS, PROC, SERPINC1*) (Germain et al., 2015; Germain et al., 2011; Rosendaal and Reitsma, 2009). This is consistent with the efficacy of therapeutic targeting of coagulation to protect against DVT with anticoagulants (Chan et al., 2016). However, although the use of anticoagulants is effective, dosing and efficacy are limited by the increase in the risk of bleeding in treated individuals (Chan et al., 2016; Schulman et al., 2014; Schulman et al., 2009; Schulman et al., 2013; Wolberg et al., 2015). Therefore, alternative targets that inhibit DVT disease processes, but that do not modify bleeding risk may provide new adjunctive therapies to further protect against the development or recurrence of DVT. GWAS studies have also identified additional risk *loci* for VTE, but with no known role in coagulation (Apipongrat et al., 2019; Germain et al., 2015; Hinds et al., 2016). This provides encouragement that alternative therapeutic targets may exist with the potential to modify the disease process without affecting bleeding risk. These *loci* include *SLC44A2* and *TSPAN15* genes (Apipongrat et al., 2019; Germain et al., 2015; Hinds et al., 2016). Despite the identification of these *loci*, the function of these cell surface proteins with respect to their involvement in the pathogenesis of venous thrombosis remains unclear.

Using microfluidic flow channels to enable analysis of the phenotypic effects of VWF-GPIbα signaling under flow, we confirm the rapid activation of the platelet integrin, α_IIb_β_3_. Activated α_IIb_β_3_ is capable of binding directly to neutrophils via a direct interaction with SLC44A2. Under flow, this interaction transduces a signal into neutrophils capable of driving NETosis. A single nucleotide polymorphism (SNP; rs2288904-A) in *SLC44A2* (minor allele frequency 0.22) that is protective against VTE (Germain et al., 2015) encodes a R154Q substitution in the first extracellular loop of the receptor that markedly reduces neutrophil-platelet binding via activated α_IIb_β_3_. These results provide a functional explanation for the protective effects of the rs2288904-A SNP and highlight the potential of SLC44A2 as an adjunctive therapeutic target in DVT.

## Results

To explore the influence of platelet binding to VWF under flow upon platelet function, full length (FL-) human VWF was adsorbed directly onto microfluidic microchannel surfaces, or the isolated recombinant VWF A1 domain, or an A1 domain variant (Y1271C/C1272R, termed A1*) that exhibits a 10-fold higher affinity for GPIbα (**Fig S1**) (Blenner et al., 2014), were captured via their 6xHis tag. Fresh blood anticoagulated with D-phenylalanyl-prolyl-arginyl chloromethyl ketone (PPACK) and labelled with DiOC_6_, was perfused through channels at 1000s^-1^ for 3.5 minutes. On FL-VWF, A1 or A1*, a similar time-dependent increase in platelet recruitment/surface coverage was observed (**Fig 1a & Fig S2**). Platelets rolled prior to attaching more firmly on all VWF channels. However, median initial platelet rolling velocity on VWF A1 was 1.76 μms^-1^, whereas on A1* this was significantly slower (median 0.23μms^-1^) (**Fig 1b & c**), reflective of its 10-fold higher affinity for GPIbα (**Movie 1**)

**Figure 1:**
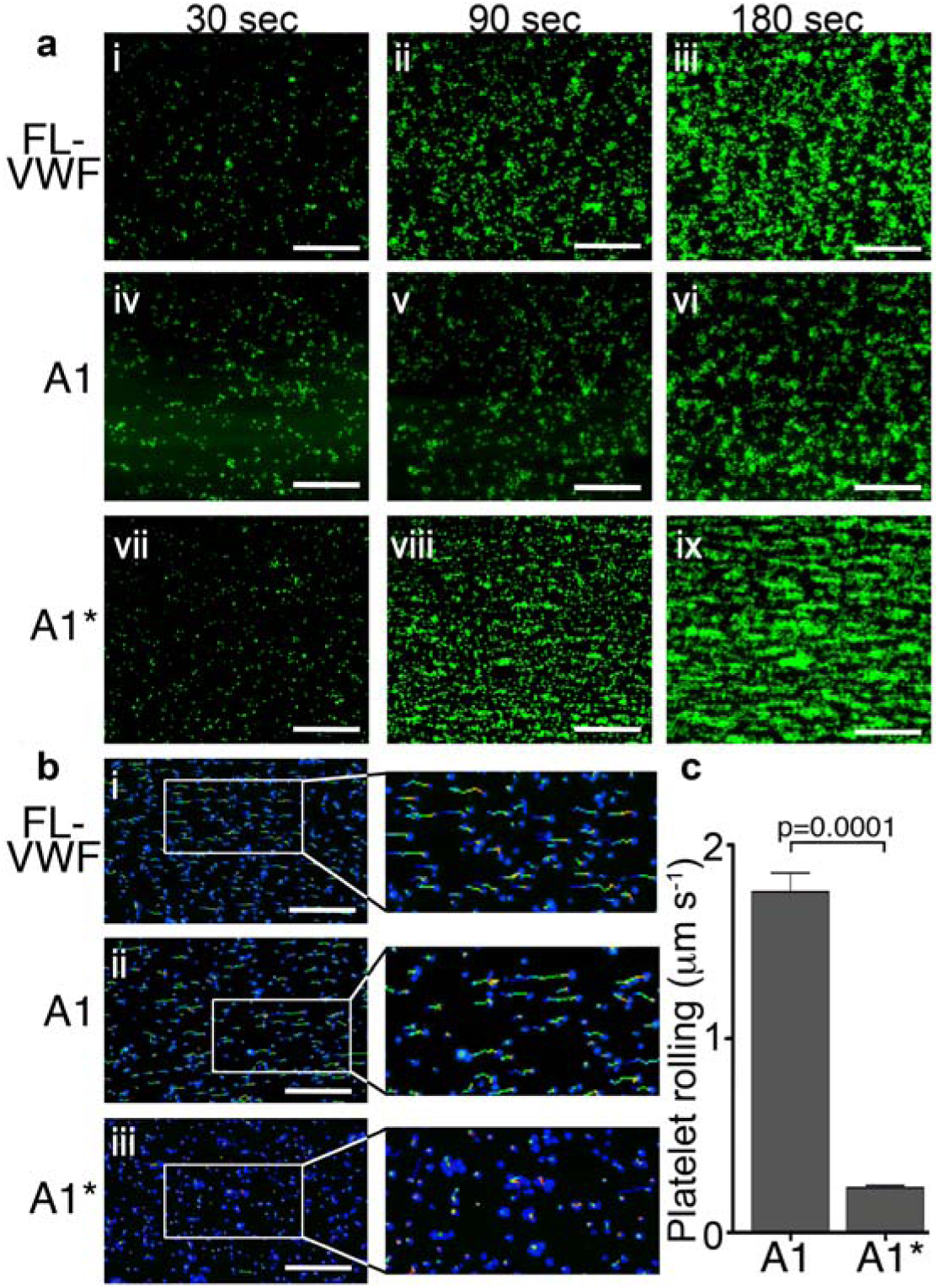
Platelet rolling and attachment to VWF under flow. **a)** Vena8 microchannels were coated with either full-length VWF (FL-VWF; i-iii), VWF A1 (iv-vi) or A1* (vii-ix). Whole blood labelled with DiOC_6_ was perfused at 1000s^-1^. Representative images (n=3) of platelets (green) after 30, 90 and 180 seconds are shown. Scale; 50μm (see also **Movie 1**). **b)** Experiments performed as in a), bound platelets (blue) were tracked (depicted by multi-coloured lines) representing distance travelled in the first 30 sec of flow. Scale bar; 50μm. **c)** Platelet rolling velocity on channels coated with A1 and A1*. Data plotted are median ±95% CI. n=3562 platelets from 3 different experiments (A1) and n=4047 platelets from 3 different experiments (A1*). Data were analyzed using the Mann-Whitney test.

### Platelet binding to VWF under flow induces intraplatelet signaling and activation of α_IIb_β_3_

Platelets bound to either FL-VWF, A1 or A1* formed small aggregates after about 2 minutes (**Fig 2ai**) due to activation of the platelet integrin, α_IIb_β_3_, and its binding to plasma fibrinogen. Consistent with this, when plasma-free blood (i.e. RBCs, leukocytes and platelets resuspended in plasma-free buffer) was used, platelets remained as a uniform monolayer, and did not form microaggregates (**Fig 2aii**). Similarly, when activated α_IIb_β_3_ was blocked in whole blood with eptifibatide or GR144053, aggregation was also inhibited (**Fig 2aiii & iv**). Irrespective of the surface (VWF, A1 or A1*), platelet aggregation was markedly reduced if plasma-free blood was used, or if α_IIb_β_3_ was blocked (**Fig 2b-e**). These results demonstrate that the A1-GPIbα interaction leads to activation of α_IIb_β_3_, which is consistent with previous reports (Goto et al., 1995; Kasirer-Friede et al.). In support of this, fluorescent fibrinogen bound to platelets tethered via FL-VWF, but not to platelets captured to channel surfaces using an anti-PECAM-1 antibody (**Fig S3a**).

**Figure 2:**
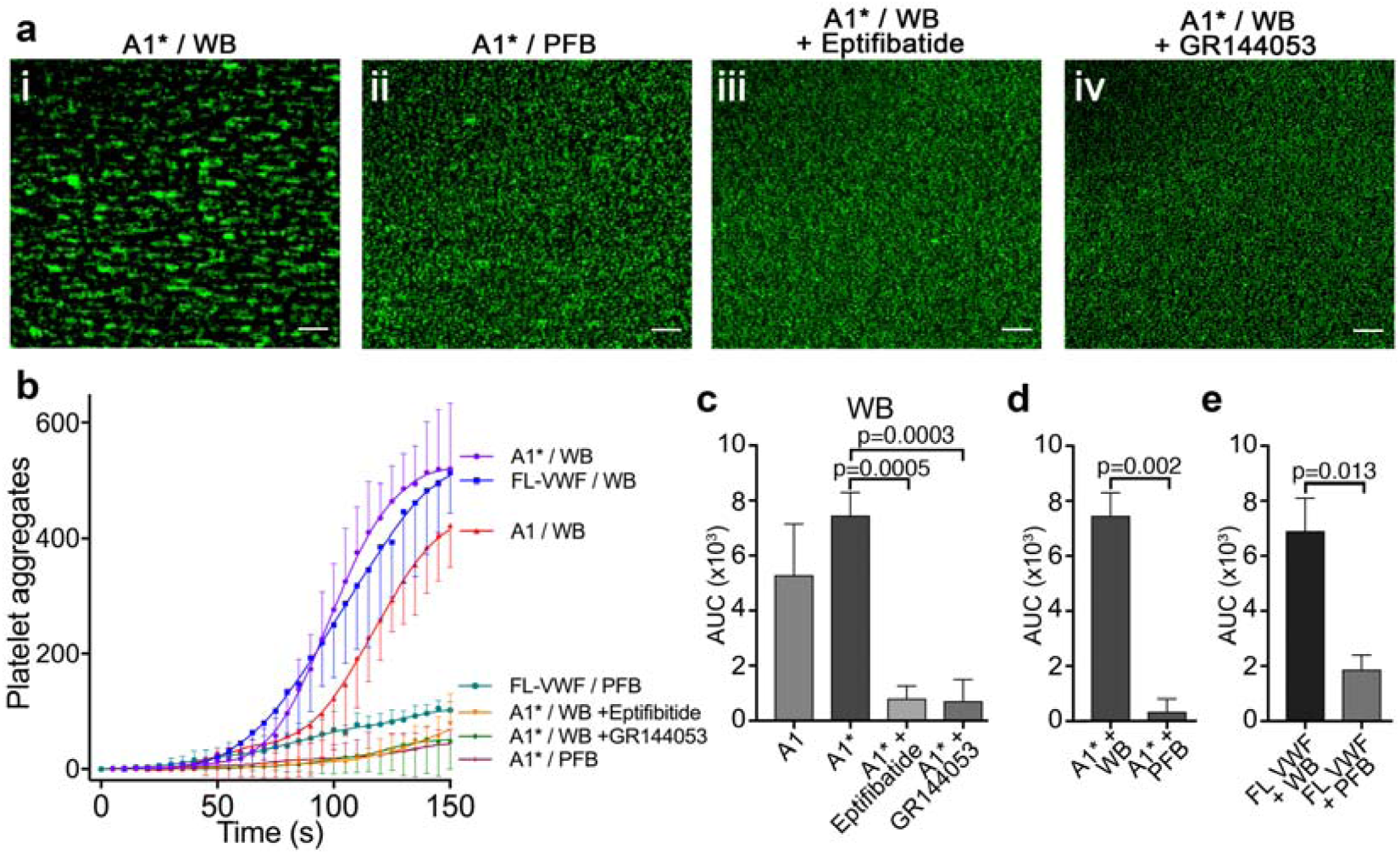
Platelet binding to VWF under flow induces α_IIb_β_3_-dependent aggregation. **a)** Vena8 microchannels were coated with A1* via its 6xHis tag. i) Whole blood (WB) or ii) plasma-free blood (PFB), iii) WB containing eptifibatide or iv) WB containing GR144053 were perfused through channels at 1000s^-1^. Representative images acquired after 3 minutes. Scale bar; 50μm. **b)** Graph measuring platelet aggregation over time in WB perfused through channels coated with A1 (red, n=3), A1* (purple, n=4) and FL-VWF (blue, n=3), WB pre-incubated with eptifibatide (orange, n=3) or GR144053 (green, n=4) over channels coated with A1* and PFB over channels coated with A1* (magenta, n=3) or FL-VWF (teal, n=3). Data plotted are mean ±SD. **c-e)** Bar charts comparing area under the curve (AUC) of the data presented in b). **c)** WB perfused over A1 or A1* with or without eptifibatide or GR144053. **d)** WB or PFB perfused over A1*. **e)** WB or PFB perfused over FL VWF. Data presented are mean ±SD, n=3 or 4 as indicated in b). Data were analyzed using the Mann-Whitney test.

To investigate the effect of A1-GPIbα-dependent signaling, platelets were preloaded with the Ca^2+^-sensitive fluorophore, Fluo-4 AM. Platelets bound to A1* under flow exhibited repeated transient increases in fluorescence, corresponding to Ca^2+^ release from platelet intracellular stores in response to A1-GPIbα binding under flow (**Movie 2**) (Kasirer-Friede et al., 2004; Mu et al., 2010). Despite intracellular Ca^2+^ release, this did not lead to appreciable P-selectin exposure (i.e. α-granule release) (**Fig S3b**). Intraplatelet Ca^2+^ release was not detected when platelets were captured under flow using an anti-PECAM1 antibody. We therefore propose that flow-dependent VWF-GPIbα signaling ‘primes’, rather than activates, platelets. This priming is characterized by activation of α_IIb_β_3_, but minimal α-granule release, and represents part of the tunable response of platelets.

### Platelets ‘primed’ by VWF interact with leukocytes

To explore the influence of platelet ‘priming’ upon their ability to interact with leukocytes, platelets were captured and ‘primed’ on VWF for 3 minutes at 1000s^-1^. Thereafter, leukocytes in whole blood (also labelled with DiOC_6_) were perfused at 50s^-1^ and rolled on the platelet-covered surface (**Movie 3 & Fig S4a**). Leukocytes did not interact with platelets captured via an anti-PECAM-1 antibody (**Fig 3a**), demonstrating the dependency on prior A1-GPIbα-mediated platelet ‘priming’.

**Figure 3:**
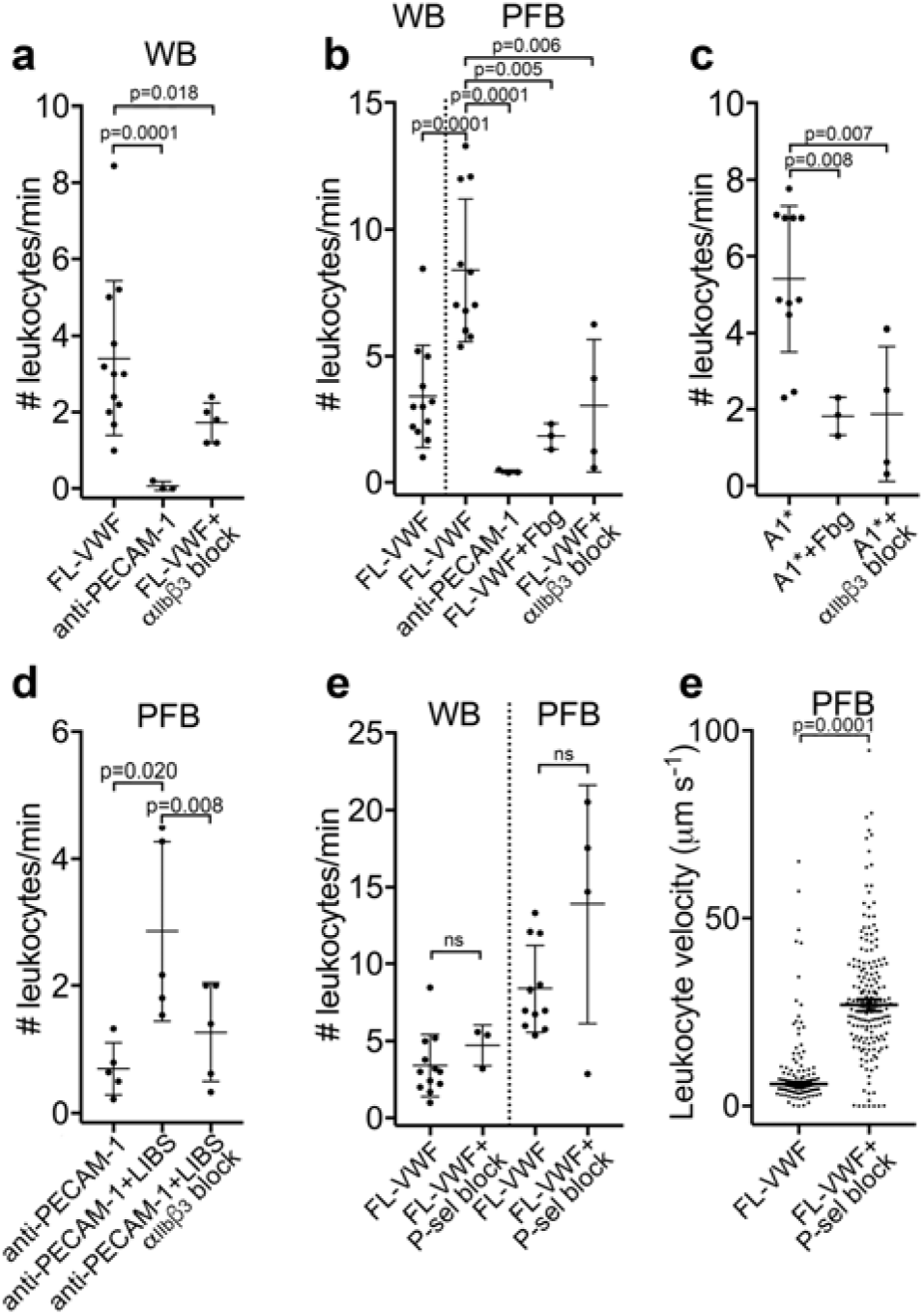
Leukocytes bind to VWF-bound platelets under flow (see also Movie 3). **a)** Graph of the number of leukocytes/minute in WB interacting with platelets bound to FL-VWF in the absence (n=12) or presence of eptifibatide/GR144053 (n=5), or binding to platelets bound to anti-PECAM-1 antibody (n=3). **b)** Graph of the number of leukocytes/minute in WB or PFB interacting with platelets bound to FL-VWF in the absence (n=12) or presence of 1.3mg/ml fibrinogen (n=3) or eptifibatide/GR144053 (n=4), or binding to platelets bound to anti-PECAM-1 antibody (n=3). **c)** Graph of the number of leukocytes/minute in PFB interacting with platelets bound to A1* in the absence (n=11) or presence of fibrinogen (n=3) or eptifibatide/GR144053 (n=4). **d)** Graph of the number of leukocytes/minute in PFB interacting with platelets bound to anti-PECAM-1 antibody in the absence (n=5) or presence of LIBS/anti-β_3_ activating antibody (n=5) ± GR144053 (n=5). **e)** Graph of the number of leukocytes/minute in WB or PFB, as shown, interacting with platelets bound to FL-VWF in the absence or presence of a blocking anti-P-selectin antibody. **f)** Graph of leukocyte rolling velocity on platelets bound to FL-VWF in PFB in the absence or presence of a blocking anti-P-selectin antibody. Data shown are individual leukocyte rolling velocities (n=121 and 178, respectively) for 3 separate experiments. In all graphs, data plotted are mean ±SD. Data were analyzed using unpaired, two-tailed Student’s t test; ns not significant.

As VWF-’primed’ platelets present activated α_IIb_β_3_, we hypothesized that ‘outside-in’ integrin signaling might be important for platelet-leukocyte interactions to occur (Durrant et al., 2017). Contrary to this, we observed a significant (∼2-fold) increase in the number of leukocytes interacting with the VWF-bound platelets in plasma-free conditions (**Fig 3b**). Moreover, addition of purified fibrinogen to plasma-free blood to 50% normal plasma concentration significantly reduced platelet-leukocyte interactions (**Fig 3b**) suggesting that leukocytes and fibrinogen compete for binding ‘primed’ platelets. Blocking α_IIb_β_3_ (**Fig 3a-b & Fig S4a**) also significantly decreased platelet-leukocyte interactions irrespective of whether platelets were captured on FL-VWF or A1*, or whether experiments were performed in whole blood or plasma-free blood (**Fig 3a-c)**.

To explore the role of activated α_IIb_β_3_ in binding leukocytes, platelets were captured onto anti-PECAM-1 coated channels and an anti-β_3_ antibody (ligand induced binding site – LIBS) that induces activation of α_IIb_β_3_ applied (Du et al., 1993). Antibody-mediated activation of α_IIb_β_3_ caused a significant increase in the number of leukocytes binding in a manner that could be blocked with GR144053 (**Fig 3d**).

The best characterized platelet-leukocyte interaction is mediated by P-selectin on activated platelets binding to PSGL-1 on leukocytes (Vandendries et al., 2004). Although we detected little/no P-selectin on the surface of VWF-’primed’ platelets, this did not formally exclude a role for P-selectin in leukocyte adhesion. Therefore, we first established the efficacy of P-selectin blockade through the marked reduction of leukocyte binding to collagen captured/activated platelets (**Fig S4b-c**). However, blockade of P-selectin on FL-VWF-bound platelets from whole blood or plasma-free blood had no effect upon the number of leukocytes interacting with the platelet surface, suggesting that the recruitment of leukocytes is independent of P-selectin (**Fig 3e**). Leukocytes rolled faster over platelet surfaces after blocking P-selectin in plasma-free blood (**Fig 3f & Movie 3**) or whole blood (**Fig S4d**), suggesting that whereas leukocyte capture is highly dependent on activated α_IIb_β_3_ (and not P-selectin), once recruited, small amounts of P-selectin on the platelet surface may slow leukocyte rolling.

### Leukocytes bind directly to activated α_IIb_β_3_

To more specifically test the leukocyte interaction with activated α_IIb_β_3_ (and to exclude other platelet receptors), purified α_IIb_β_3_ was covalently coupled to microchannels and, thereafter, activated with Mn^2+^ (Litvinov et al., 2005). Isolated PBMCs and PMNs were perfused through α_IIb_β_3_-coated channels at 50s^-1^. Cells from both PBMCs and PMNs (**Fig 4ai-ii & 4b**) directly attached to the activated α_IIb_β_3_ surface. This binding was significantly diminished (>70%) by adding either purified fibrinogen or eptifibatide to the PMNs (**Fig 4aiv-v & 4b**).

**Figure 4:**
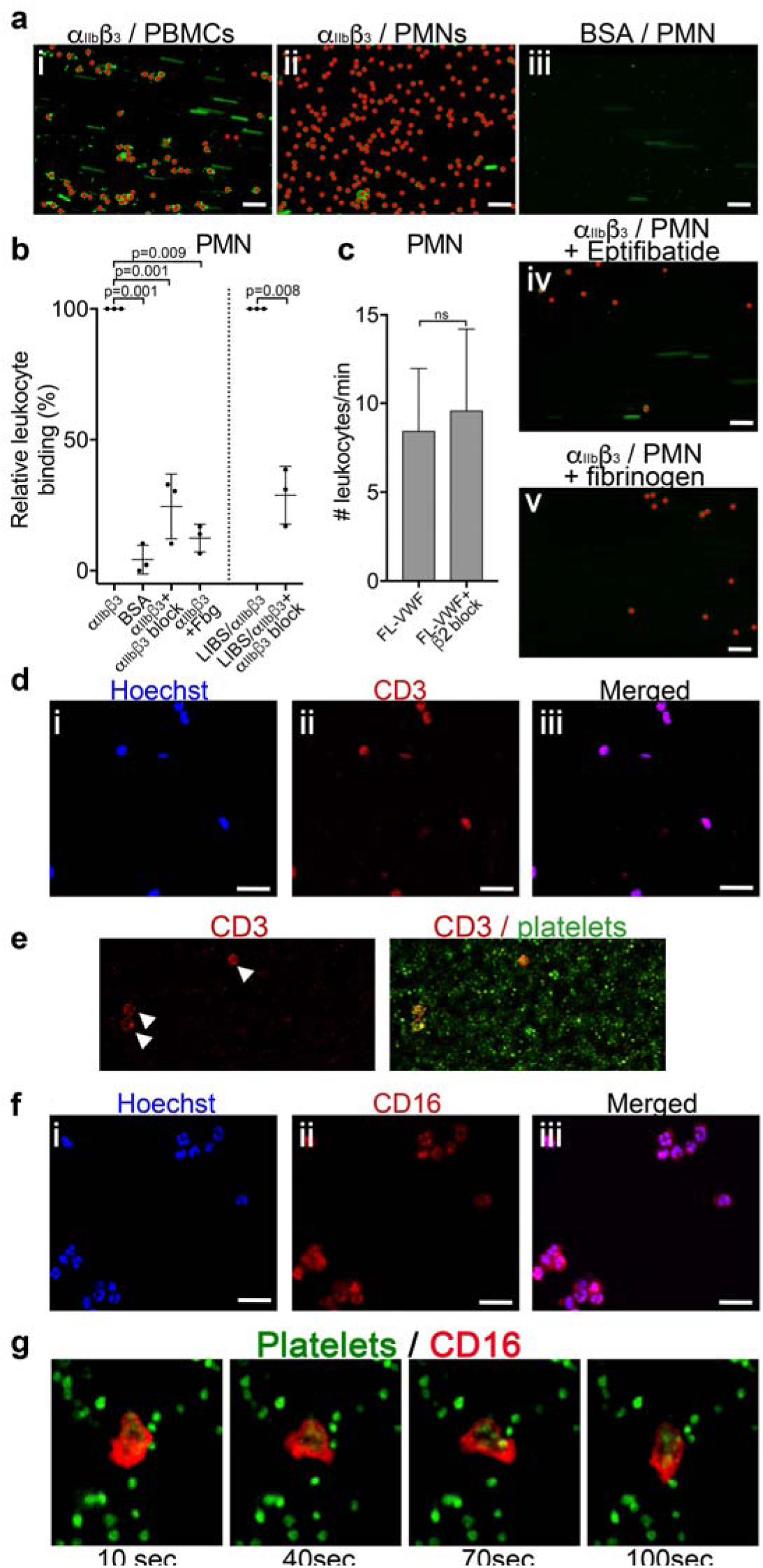
Leukocytes bind to activated α_IIb_βunder flow. **a)** Purified α_IIb_β_3_ or BSA, as noted, were covalently coupled to microchannel surfaces by amine coupling. α_IIb_β_3_ was activated using Mn^2+^ and Ca^2+^ in all buffers. PBMCs (i) or PMNs (ii-v) labelled with DiOC_6_ were perfused through channels at 50s^-1^ in the presence and absence of eptifibatide (iv) or 1.3mg/ml purified fibrinogen (v). Bound leukocytes (as opposed to flowing) are pseudo-colored red to aid visualization and to distinguish from leukocytes in transit. Scale bar; 50μm. **b)** Graphical representation of relative leukocyte binding to activated α_IIb_β_3_ in the presence and absence of eptifibatide or 1.3mg/ml purified fibrinogen, or to BSA after 15 minutes of PMN perfusion (n=3), or to α_IIb_β_3_ captured and activated by LIBS/anti-β_3_ activating antibody in the absence and presence of GR144053 (n=3). Data plotted are mean ±SD. Data were analyzed using unpaired, two-tailed Student’s t test. **c)** Graph of the number of leukocytes/minute in PFB interacting with platelets bound to FL-VWF in the absence (n=12) or presence of a blocking anti-β2 integrin polyclonal antibody (n=5) capable of blocking both LFA-1 or Mac-1 on leukocytes. **d)** PBMCs stained with Hoechst dye (i - blue), anti-CD3 (ii - red) and merged (iii). Representative of n=4. Scale bar; 20μm. **e)** PFB stained with DiOC6 was perfused over FL-VWF at 1000s^-1^ followed by 50s^-1^. T-cells labelled with anti-CD3 (red - arrows) were seen to attach to ‘primed’ platelets **f)** PMNs stained with Hoechst dye (i - blue), anti-CD16 (ii - red) and merged (iii). Representative of n=4. Scale bar; 20μm (see also **Movie 4**). **g)** Images depicting a neutrophil stained with anti-CD16 (red) ‘scanning’ the ‘primed’ platelets stained with DiOC_6_ (green). Images shown were taken 10, 40, 70 and 100 seconds after neutrophil attachment note the movement of the neutrophil shown - see also **Movie 4**.

We also captured purified α_IIb_β_3_ to flow channel surfaces using the activating anti-β_3_ (LIBS) antibody. Leukocytes were again efficiently captured to this surface in a manner that could be inhibited (∼70%) by blocking α_IIb_β_3_ (**Fig 4b**).

Activated (rather than resting) leukocytes can interact with platelets via Mac-1 (α_M_β_2_) (either directly through GPIbα or via fibrinogen bridge with activated α_IIb_β_3_) or LFA-1 (α_L_β_2_) via ICAM-2 (Damle et al., 1992; Diacovo et al., 1994; Simon et al., 2000; Weber and Springer, 1997). However, blocking β_2_ suggested no role for either of these activated integrins in leukocyte binding to VWF-’primed’ platelets (**Fig 4c**). In summary, we show leukocytes bind α_IIb_β_3_ directly dependent upon its RGD-binding groove, but in a manner that is independent of Mac-1 or LFA-1.

### T-cells and neutrophils interact with VWF-’primed’ platelets via activated α_IIb_β_3_

We found no evidence of either CD14^+^ monocytes or CD19^+^ B-cells in PBMCs interacting with activated α_IIb_β_3_. T-cells were the only cell type amongst the PBMCs capable of binding activated α_IIb_β_3_ or VWF-‘primed’ platelets (**Fig 4d-e**).

Using isolated PMNs, we found that cells stained with anti-CD16 bound to activated α_IIb_β_3_-coated channels and also to VWF-’primed’ platelets (**Fig 4f & 4g**). Based on multi-lobulated segmented nuclear morphology (**Fig 4f**), these cells were indicative of CD16^+^ neutrophils. Neutrophils scanned the platelet- or α_IIb_β_3_-coated surfaces (**Fig 4g & Movie 4**) suggesting that the binding of neutrophils to α_IIb_β_3_ under flow may itself initiate signaling events within neutrophils. In line with this, PMNs bound to VWF-’primed’ platelets (**Fig 5a**) or activated α_IIb_β_3_ (**Fig 5b**) surfaces exhibited similar intracellular Ca^2+^ release (**Movie 5**) that reached a maximum after 200-300 seconds (**Fig 5c-d**).

**Figure 5:**
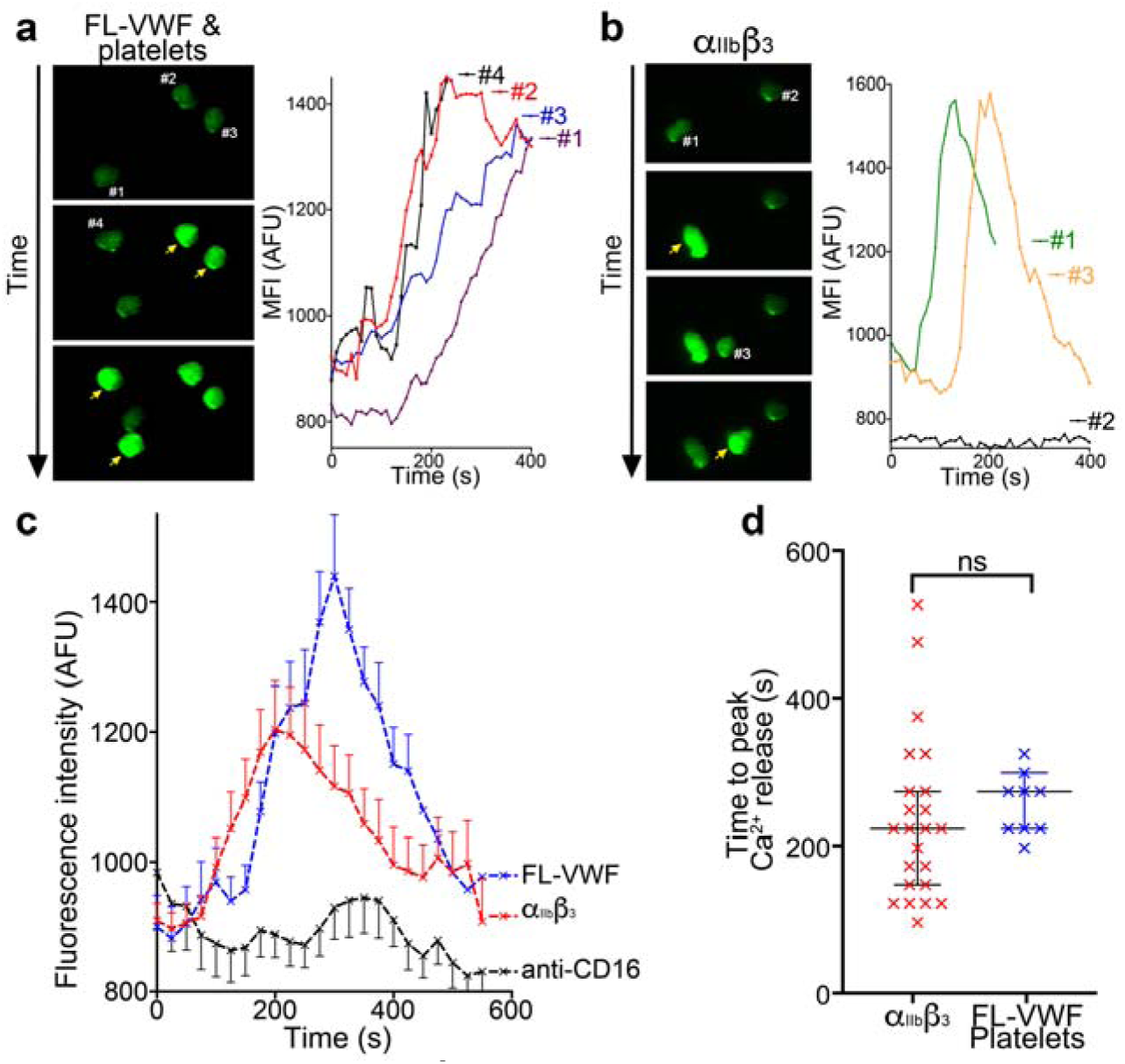
Binding to α_IIb_β_3_ induces intracellular Ca^2+^ release in neutrophils. **a)** Representative images of neutrophils pre- loaded with Fluo-4 AM bound to VWF-’primed’ platelets captured (**Movie 5**). Neutrophils are numbered #1-#4. The yellow arrow highlights a frame in which the fluorescence has increased in the attached neutrophil. For each neutrophil shown, intracellular Ca^2+^ release is quantified by measurement of cellular mean fluorescent intensity (MFI) over time. **b)** As in A) except neutrophils were perfused over activated α_IIb_β_3_. MFI increased for neutrophils #1 and #3, but not for neutrophil #2. **c)** Graph depicting the change in MFI as a function of time after neutrophil attachment to microchannels coated with activated α_IIb_β_3_ (n=24 neutrophils from 3 different experiments), VWF-’primed’ platelets (n=9 neutrophils from 1 experiment) or anti-CD16 (n=13 neutrophils from 2 different experiments). Data plotted are mean ±SEM. **d)** Dot plot presenting the time between neutrophil attachment and maximum MFI of neutrophils binding to purified α_IIb_β_3_ (red), or VWF-’primed’ platelets (blue). Data plotted are median ±95% confidence interval. Data were analyzed using the Mann-Whitney test.

### Binding of neutrophils to α_IIb_β_3_ under flow induces Nox- and Ca^2+^-dependent NETosis

Platelets assist in the targeting of intravascular bacterial pathogens through stimulation of the release of NETs (Brinkmann et al., 2004; Gaertner et al., 2017; Wong et al., 2013; Yeaman, 2014). However, the physiological agonists or mechanisms that drive NETosis are not fully resolved (Nauseef and Kubes, 2016). We therefore examined whether the binding of neutrophils to α_IIb_β_3_ might induce NETosis. Isolated PMNs were perfused over either activated α_IIb_β_3_ or anti-CD16 (negative control) at 50s^-1^ for 10 minutes and NETosis was subsequently analyzed under static conditions (**Fig 6a & Movie 6**). Nuclear decondensation was evident after ∼60 minutes, and Sytox Green fluorescence, indicative of cell permeability that precedes NETosis, was detected from ∼85 minutes. Nuclear decondensation, increased cell permeability and positive staining with a cell impermeable DNA fluorophore do not specifically identify NETosis. Therefore, anti-citrullinated histone H3 antibody was perfused through the channels after 90 minutes to more specifically identify NETs. The introduction of flow at this point caused the DNA to form extended mesh-like NETs that were stained positively by Hoechst and the anti-citrullinated histone H3 antibody (**Fig 6b**). Very similar results were obtained with PMN bound to either α_IIb_β_3_ (captured by the activating anti-β_3_ antibody), or to platelets ‘primed’ by A1* or FL-VWF. On activated α_IIb_β_3_, 69% ±14% of neutrophils through the entire channel formed NETs after 2 hours, compared to minimal NETosis events (8% ±8%) when neutrophils were captured by anti-CD16 (**Fig 6c**). When neutrophils were captured on α_IIb_β_3_ in the absence of flow, neutrophils attached, but NETosis was significantly reduced by 4-fold, with on 17% of neutrophils exhibiting signs of NETosis (**Fig 6c**). This suggested that the signaling mechanism from the platelet to the neutrophil is mechano-sensitive and does not require other platelet receptors or releasate components.

**Figure 6:**
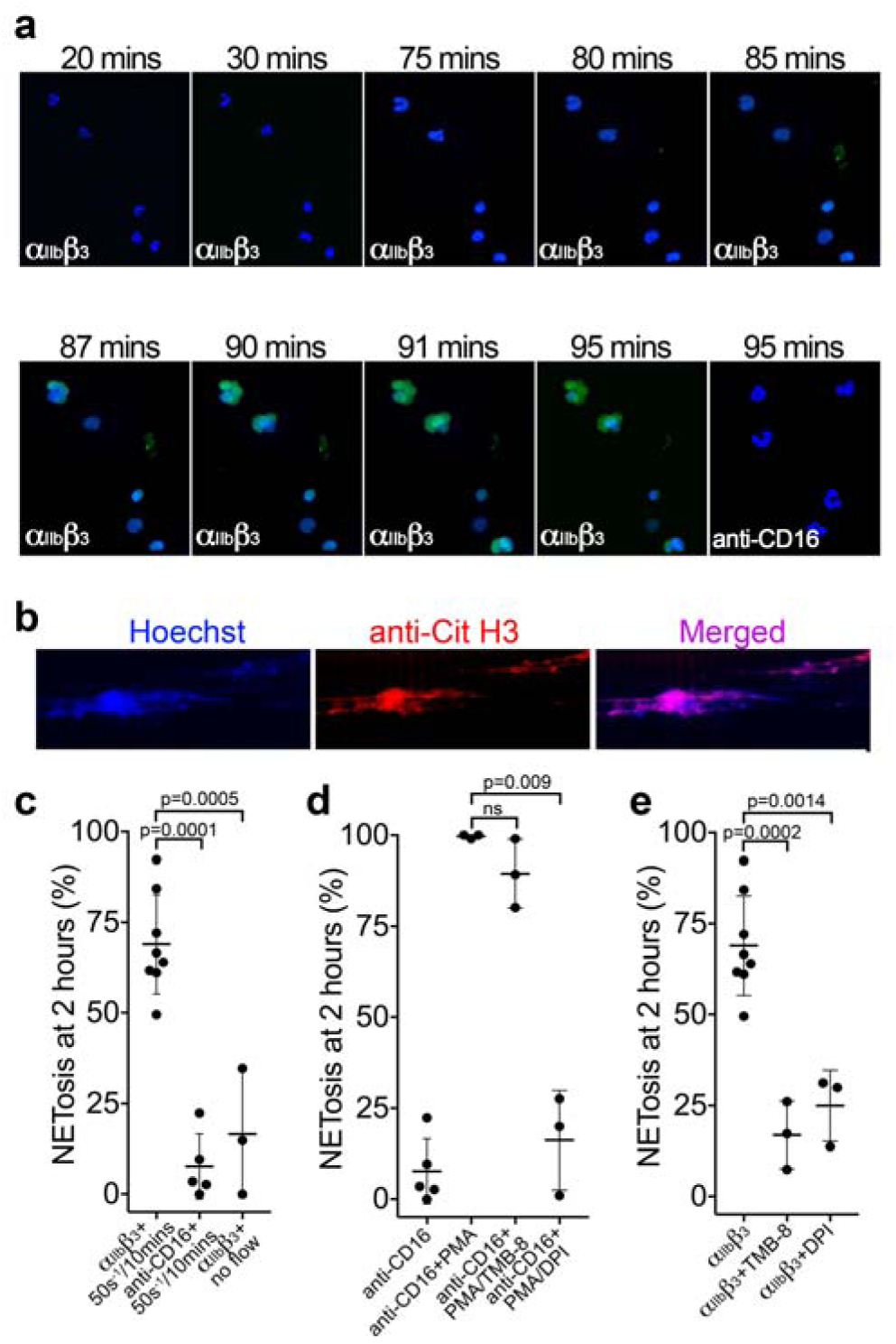
Binding of neutrophils to α_IIb_β_3_ under flow induces NETosis. **a)** Isolated PMNs labelled with Hoechst (blue) and cell-impermeable Sytox Green were perfused over α_IIb_β_3-_coated microchannels, or anti-CD16, (-ve ctrl) at 50s^-1^ for 10 minutes and then monitored under static conditions. Representative composite images after 20, 30, 75, 80, 85, 87, 90, 91 and 95 minutes of attachment. Neutrophils bound to α_IIb_β_3_ exhibited nuclear decondensation and increased cell permeability that precedes NETosis after about 85 minutes, Sytox Green staining appears, indicative of DNA becoming extracellular (see **Movie 6**). Neutrophils bound to surfaces using an anti-CD16 antibody did not exhibit signs of NETosis or did so very rarely. **b)** Immunostaining of neutrophils bound to α_IIb_β_3_ after 90 mins as in a). Hoechst (blue), citrullinated H3 (red) and merged images are shown. **c)** Graph showing the mean % of neutrophils ±SD in the entire microchannel that formed NETs after 2 hours of attachment on α_IIb_β_3_ (n=8) or anti-CD16 (n=5), captured in the presence of flow (50s^-1^/10 minutes), or captured on α_IIb_β_3_ under static/no flow conditions (n=3). **d)** Graph showing the mean % of neutrophils ±SD in the entire microchannel that formed NETs after 2 hours of attachment on anti-CD16 antibody in the presence of flow (50s^-1^/10 minutes) (n=5) in the presence of PMA (n=3), PMA and TMB-8 (n=3) or PMA and DPI (n=3). **e)** Graph showing the mean % of neutrophils ±SD in the entire microchannel that formed NETs after 2 hours of attachment on α_IIb_β_3_ in the presence of flow (50s^-1^/10 minutes) (n=8) and in the presence of TMB-8 (n=3) or DPI (n=3), as noted. Data were analyzed using unpaired, two-tailed Student’s t test; ns not significant.

Neutrophils captured by an anti-CD16 antibody and stimulated with phorbol 12-myristate 13-acetate (PMA) for two hours led to 100% ±0.5% of neutrophils releasing NETs (**Fig 6d**). PMA-induced NETosis was not significantly inhibited in the presence of TMB-8 (an antagonist of intracellular Ca^2+^ release; 90% ±9%), but was effectively inhibited in by DPI (NADPH oxidase inhibitor; 16% ±13%), similar to previous reports (Gupta et al., 2014). NETosis of neutrophils captured by α_IIb_β_3_ under flow was significantly inhibited by TMB-8 (17% ±9%) and DPI (25% ±10%) (**Fig 6e**). This highlights the dependency of both intracellular Ca^2+^ release and NADPH oxidase signaling pathways in NETosis in response to binding α_IIb_β_3_ under flow.

### ‘Primed’ platelets interact with SLC44A2 receptor on neutrophils

Our data point to the presence of a specific receptor on the surface of neutrophils (and T-cells) that is not present on B cells or monocytes and that is capable of binding to activated α_IIb_β_3_ and transducing a signal into the cell. To identify this leukocyte counter-receptor, we analyzed RNA sequencing data from different leukocyte populations, selecting genes that are expressed at higher levels in neutrophils (or in CD4^+^ T-cells) than in monocytes (Adams et al., 2012; Grassi et al., 2019) (**Fig 7**). We further limited the candidate search by selecting those genes that code for transmembrane proteins. Using this approach, we identified 93 candidate genes. Of these, 33 genes were excluded as they are primarily associated with intracellular membranes. An additional 16 genes were also excluded due to the presence of short extracellular regions/domains (<30 a.a.) that would unlikely be capable of facilitating interactions with an extracellular binding partner. (**Fig 7**) We then analyzed proteomic data to verify the preferential expression of the remaining candidates in neutrophils as opposed to monocytes (Rieckmann et al., 2017). These data suggested that the protein product of 14 of the remaining genes appeared to be detected in higher abundance in monocytes, which we used as a further exclusion criterion (**Fig S5a**). From the remaining 30 candidate genes, the *SLC44A2* gene was selected for validation due to its recent identification as a risk locus for both DVT and stroke (Germain et al., 2015; Hinds et al., 2016), both of which are pathologies associated with described contributions of platelet-leukocyte interactions. SLC44A2 is a cell surface receptor with 10 membrane-spanning domains and five extracellular loops of 178a.a., 38a.a., 72a.a., 38a.a. and 18a.a. in length, respectively (Nair et al., 2016). We sourced antibodies against SLC44A2 that specifically recognize amino acid sequences within the first and second extracellular loops. Published proteomic profiling confirmed the preferential expression of SLC44A2 in neutrophils (**Fig S5a**) (Rieckmann et al., 2017). Western blotting of isolated granulocyte lysates revealed two bands representing SLC44A2 (glycosylated and nascent/non-glycosylated SLC44A2) (**Fig S5b**). Perfusing human neutrophils over immobilized activated α_IIb_β_3_ in the presence of the first anti-SLC44A2 antibody (anti-SLC44A2 #1) that recognizes the second extracellular loop revealed a dose-dependent blockade of neutrophil binding when compared to no antibody or control rabbit IgG (**Fig 8a-b**). A second anti-SLC44A2 antibody (anti-SLC44A2 #2) that recognizes the first extracellular loop region of SLC44A2 confirmed these findings (**Fig 8a**). The anti-SLC44A2 #2 almost completely blocked neutrophil binding to activated α_IIb_β_3_ suggesting that this antibody more effectively blocks the neutrophil binding to the integrin than anti-SLC44A2 #1. This may suggest that the first and longest extracellular loop is involved in interaction with an extracellular ligand

**Figure 7:**
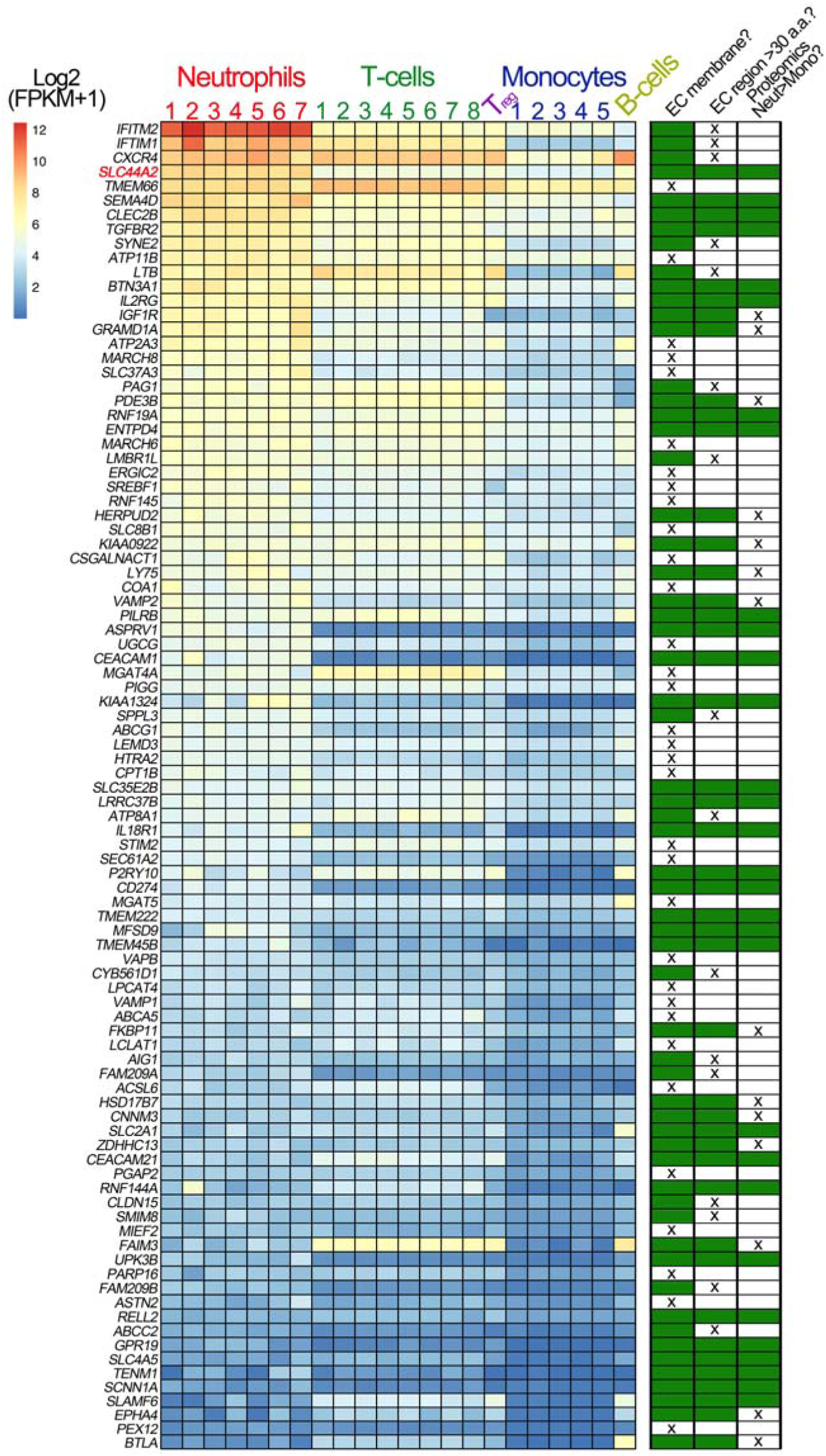
Transcriptomic profiling of human leukocytes. RNA sequencing data from different leukocytes were obtained from the BLUEPRINT consortium [https://www.biorxiv.org/content/10.1101/764613v1]. Differential gene expression analyses were performed: mature neutrophils (n=7) vs monocytes (n=5) and CD4-positive/αβ T cells (n=8) vs monocytes (n=5). Regulatory T cells (T_reg_, n=1) and native B cells (n=1), are included in the heatmap, for comparison but were not used in differential gene expression analysis due to the low number of biological replicates. We first selected genes that were expressed significantly higher in neutrophils than in monocytes, and also those that were significantly higher in CD4-positive/αβ T cells than in monocytes. Their intersection identified 750 genes (598 of which protein coding). From these 598 genes, we selected the 93 genes that contained the Uniprot annotation of “INTRAMEMBRANE DOMAIN” or “TRANSMEM DOMAIN”. The effective log2(FPKM+1) data are presented in the heatmap of the 93 genes, with the rows ordered according to the mean neutrophil expression levels. Next to the heatmap is a table highlighting the subsequent selection criteria used to further narrow the search for candidate receptors for α_IIb_β_3_. The first round of selection involved discarding those transmembrane proteins that are not present on the extracellular membrane, or primarily associated with intracellular membranes. The second selection criterion was to discard those proteins that had extracellular regions of <30 amino acids that might be less likely capable of mediating specific ligand binding. Finally, analysis of proteomic data from the ImmProt (http://immprot.org) resource was used to verify higher levels of protein of each selected gene in neutrophils than in monocytes

**Figure 8:**
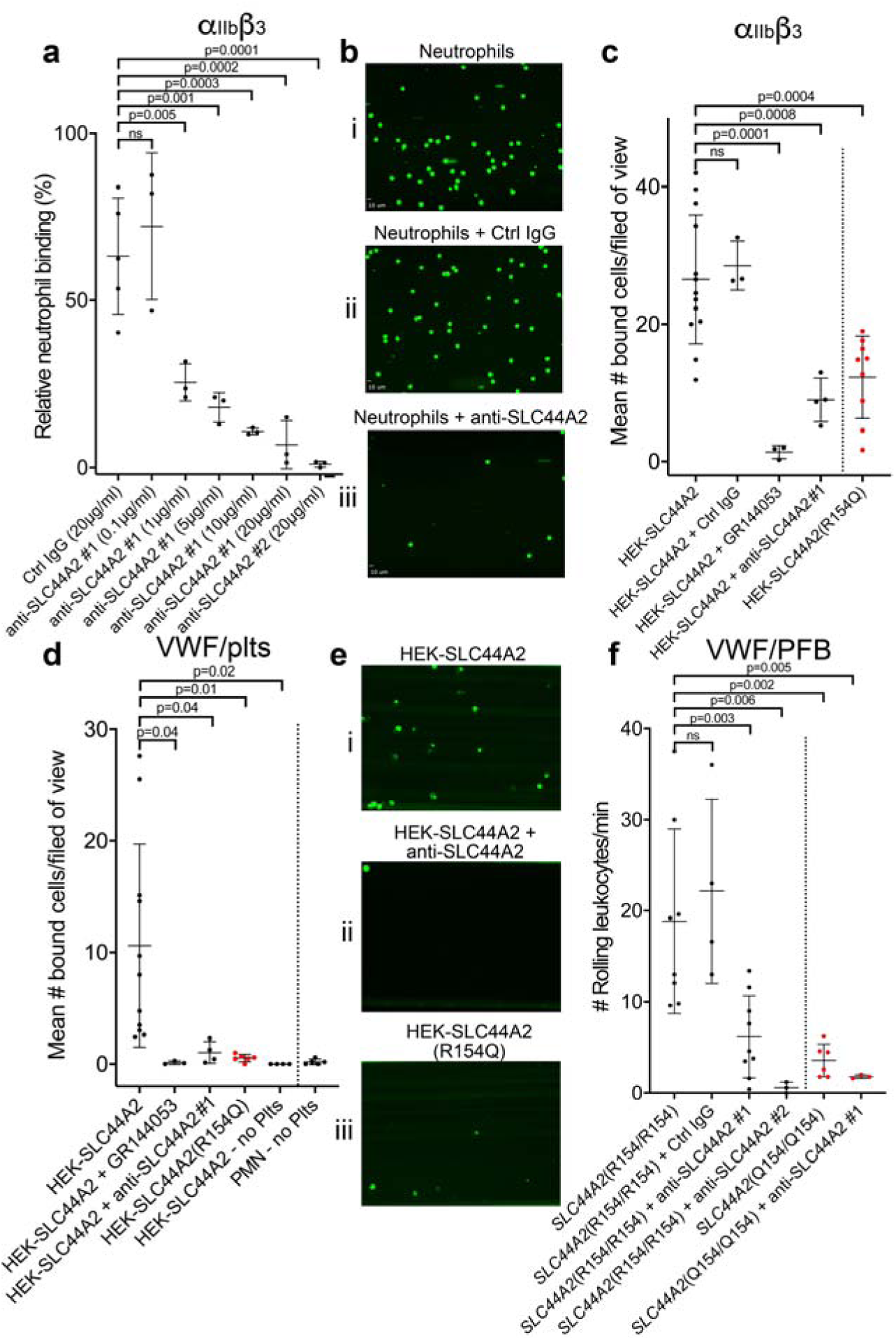
SLC44A2 binds activated α_IIb_β_3_. **a)** Graphical representation of relative neutrophil binding after 15 minutes of leukocyte perfusion at 50s^-1^ to activated α_IIb_β_3_ captured and activated by LIBS2/anti-β_3_ activating antibody in the presence and absence of increasing concentrations of anti-SLC44A2#1 or #2 antibodies. Data plotted are mean ±SD. Data were analyzed using One-Way ANOVA with multiple comparisons. **b)** Representative micrographs of neutrophils bound to activated α_IIb_β_3_ i) in the absence of antibody, ii) in the presence of control IgG and iii) in the presence of anti-SLC44A2#1. **c)** Graphical representation of the number of HEK293T cells transfected with either SLC44A2-EGFP or SLC44A2(R154Q)-EGFP (shown in red) expression vectors binding to activated α_IIb_β_3_ (captured and activated by LIBS2/anti-β3 activating antibody) after 10 mins flow at 25s^-1^. Experiments were performed in the presence and absence of either GR144053 or anti-SLC44A2#1 antibody. Data presented are the mean number of bound cells per field of view. Data were analyzed using One-Way ANOVA with multiple comparisons. **d)** Graphical representation of the number of HEK293T cells transfected with either SLC44A2-EGFP or SLC44A2(R154Q)-EGFP (shown in red) expression vectors interacting with VWF-’primed’ platelets. Plasma-free blood was first perfused at 1000s^-1^ for 3.5 mins to capture and ‘prime’ the platelets, and transfected HEK293T cells were subsequently perfused at 25s^-1^ for 10 mins, in the presence and absence of GR144053 or anti-SLC44A2#1 antibody. HEK293T cells transfected with SLC44A2-EGFP or isolated neutrophils were also perfused over VWF in the absence of platelets, for 30 mins at 25s^-1^ or 50s^-1^ respectively. Data presented are the mean number of bound cells per field of view. Data were analyzed using One-Way ANOVA with multiple comparisons. **e)** Representative micrographs of HEK293T cells transfected with i) SLC44A2-EGFP in the absence of antibody, ii) SLC44A20EGFP in the presence of anti-SLC44A2#1 antibody and iii) SLC44A2(R154Q)-EGFP bound to activated VWF-’primed’ platelets. **f)** Graphical representation of the number of neutrophils rolling per minute on VWF-’primed’ platelets. PFB from individuals homozygous for the R154-encoding allele of *SLC44A2* (R154/R154) or the Q154-encoding allele of *SLC44A2* (Q154/Q154) (shown in red) were perfused over ‘primed’ platelets for 10 mins at 50s^-1^ (see **Movie 7**). Experiments were performed in the presence and absence of anti-SLC44A2#1 or anti-SLC44A2#2 antibodies. Data plotted are mean ±SD. Data were analyzed using One-Way ANOVA with multiple comparisons.

Based on these results, we transfected HEK293T cells with an expression vector for human SLC44A2 fused to EGFP at the intracellular C-terminus. Transfected cells were perfused through activated α_IIb_β_3_ coated channels and cell binding was quantified. Transfected cells bound to these surfaces in a manner that could be blocked by GR144053 (that blocks α_IIb_β_3_) or by the anti-SLC44A2 #1 antibody (**Fig 8c**).

The SNP in *SLC44A2* identified by GWAS studies that is protective against VTE and stroke (rs2288904-A) causes a missense mutation (R154Q) in the first 178a.a. extracellular loop of SLC44A2 (Germain et al., 2015). Based on this, we hypothesized that this substitution might exert a functional influence upon the ability of SLC44A2 to interact with α_IIb_β_3_. Consistent with this hypothesis, HEK293T cells transfected with the SLC44A2 (R154Q)-EGFP expression vector exhibited reduced ability to interact with immobilized α_IIb_β_3_ (**Fig 8c**).

To further explore the potential interaction between SLC44A2 and activated α_IIb_β_3_ on platelets, we first captured and ‘primed’ platelets over VWF-coated surfaces and, thereafter, perfused SLC44A2-EGFP-transfected HEK293T cells. Again, these cells bound to VWF-’primed’ platelets in a manner that could be blocked completely with GR144053 (to block α_IIb_β_3_) or the anti-SLC44A2 #1 antibody (**Fig 8d-e**). Consistent with the previous results, HEK293T cells transfected with SLC44A2(R154Q) exhibited markedly reduced binding to VWF-’primed’ platelets (**Fig 8d-e**). A previous report suggested that SLC44A2 might bind directly to VWF (Bayat et al., 2015). However, when SLC44A2-EGFP-transfected HEK293T cells were perfused of VWF surfaces, in the absence of platelets, no binding was detected (**Fig 8d**). Similarly, isolated neutrophils also failed to interact directly with VWF coated surfaces, demonstrating the absolute dependence of platelets in facilitating cell capture under flow.

### Neutrophils homozygous for the rs2288904-A SNP exhibit reduced binding to activated α_IIb_β_3_

The rs2288904-A SNP in *SLC44A2* has a minor allele frequency of 0.22 and is protective against VTE (Germain et al., 2015). It is therefore the common allele, rs2288904-G, that is the risk allele for VTE with an odds ratio of 1.2-1.3. The frequency of individuals homozygous for the protective rs2288904-A allele amongst VTE cases is 30%-50% lower than in healthy controls. Given its prevalence, we genotyped a group of healthy volunteers to identify individuals homozygous for the major allele (rs2288904-G/G), *SLC44A2* (R154/R154), and for the protective allele (rs2288904-A/A), SLC44A2 (Q154/Q154) (**Fig S5c**). *SLC44A2* (R154/R154) neutrophils interacted with VWF-’primed’ platelets as before (**Fig 8f & Movie 7**). Consistent with the previous blocking experiments, this binding was partially blocked with anti-SLC44A2 #1, and almost completely blocked by anti-SLC44A2 #2 (**Fig 8f & Movie 7**). Furthermore, and consistent with the transfection studies, neutrophils homozygous for the protective allele, *SLC44A2* (Q154/Q154), exhibited markedly reduced (∼75%) binding to VWF-’primed’ platelets (**Fig 8f & Movie 7**) demonstrating a functional consequence of the rs2288904-A polymorphism on this neutrophil-platelet interaction.

## Discussion

Although the ability of platelet GPIbα binding to VWF to mediate intraplatelet signaling events has been known for many years, the role that it fulfils remains poorly understood (Goto et al., 1995). We demonstrate that under flow GPIbα-A1 binding ‘primes’, rather than activates, platelets, based on the rapid activation of α_IIb_β_3_, but the lack of appreciable surface P-selectin exposure (**Fig 2 & Fig S3**). Some studies have reported that GPIbα-VWF-mediated signaling can induce modest α-granule release. However, the use of static conditions and processing of platelets may explain those observations. Despite this, when compared to other platelet agonists, degranulation and P-selectin exposure induced by VWF binding are both very low (de Witt et al., 2014; Deng et al., 2016).

That platelet binding to VWF under flow ‘primes’, rather than activates, platelets is consistent with *in vivo* observations. At sites of vessel damage, VWF is important for platelet accumulation through all layers of the hemostatic plug (Joglekar et al., 2013; Lei et al., 2014; Verhenne et al., 2015). All platelets within a thrombus/hemostatic plug likely form interactions with VWF. Despite this, it is only the platelets in the ‘core’ of the thrombus that become P-selectin-positive, procoagulant platelets, whereas the more loosely bound platelets that form the surrounding ‘shell’ remain essentially P-selectin-negative (Welsh et al., 2014). If VWF-binding alone were sufficient to fully activate platelets, the differential platelet characteristics of the ‘core’ and ‘shell’ would not be observed.

Although it is frequently implied that VWF is only important for platelet capture under high shear conditions, murine models of venous thrombosis with no collagen exposure have repeatedly revealed an important role for VWF-mediated platelet accumulation (Bergmeier et al., 2008; Brill et al., 2011; Chauhan et al., 2007). Platelet binding to VWF occurs most efficiently at arterial shear rates, but still occurs under lower linear venous shear (Miyata and Ruggeri, 1999; Yago et al., 2008; Zheng et al., 2015). However, linear channels do not mimic the distorted and branched paths of the vascular system that cause more disturbed flow patterns, particularly around valves. Using channels with changing geometry under lower shear conditions, we and others have noted that VWF captures platelets appreciably more efficiently in areas of disturbed flow (Zheng et al., 2015). Indeed at venous flow rates through bifurcated channels (**Fig S6a**), we detected platelet capture on FL-VWF with concomitant ‘priming’ and leukocyte binding (**Fig S6b**). This was appreciably augmented at bifurcation points where disturbed flow exists. Consistent with our earlier findings, leukocyte binding was almost completely inhibited when α_IIb_β_3_ was blocked (**Fig S6c**). This implies that VWF can function in platelet recruitment within the venous system, particularly in areas of turbulence (e.g. branch sites, valves), which are frequently the nidus for thrombus formation in DVT.

In venous thrombosis, the thrombus generally forms over the intact endothelium, in the absence of vessel damage. This poses the question of how VWF might contribute to DVT if subendothelial collagen is not exposed. It is likely that this reflects the function of newly-secreted ultra-large VWF released from endothelial cells. Under low disturbed flow, released ultra-large VWF may tangle to form strings/cables over the surface of the endothelium. Tangled VWF strings/cables are appreciably more resistant to ADAMTS13 proteolysis than VWF that is simply unraveled. In the murine stenosis model of DVT, complete VWF-deficiency prevents platelet binding over the endothelium (Bergmeier et al., 2008; Brill et al., 2011; Chauhan et al., 2007). Similarly, blocking GPIbα binding to VWF also completely blocks platelet accumulation and thrombus formation in the stenosis model of DVT. Thus, when platelets bind to VWF under flow in such settings, platelets may become ‘primed’ facilitating both aggregation and neutrophil binding through activated α_IIb_β_3_, but without activating them into procoagulant platelets.

Our study reveals, that T-cells and neutrophils can bind directly to activated α_IIb_β_3_ on platelets and that is coupled to microchannel surfaces (**Fig 3a-d & Fig 4a-b**). In both cases, the interaction was inhibited by eptifibatide and GR144053 suggesting that both cell types may share the same receptor, in a manner that is dependent upon the RGD binding groove of activated α_IIb_β_3_.

Previous studies have identified roles for β_2_ integrins, Mac-1 (α_M_β_2_) and LFA-1 (α_L_β_2_), on leukocytes in mediating interactions with platelets. It should be recognized that the interactions of these molecules are dependent upon the integrins first being activated (and therefore also the cell), which is not the case in our system and is in contrast to previous studies implicating Mac-1 and LFA-1. However, we provide evidence that Mac-1 (α_M_β_2_) and LFA-1 (α_L_β_2_) are not involved by: 1) Leukocytes do not bind to ‘unprimed’ platelets captured by anti-PECAM-1. As Mac-1 and LFA-1 bind to GPIbα and ICAM-2, respectively, both of which are constitutively presented on the platelet surface (Kuijper et al., 1998; Simon et al., 2000), if Mac-1 and LFA-1 were the receptors involved binding would have been observed in these experiments (**Fig 3a-b**). 2) Mac-1 on leukocytes can bind indirectly to activated αIIbβ3 via a fibrinogen bridge (Weber and Springer, 1997). However, we demonstrate that fibrinogen competes for leukocyte binding to bind to activated αIIbβ3 (**Fig 3b-c & Fig 4b**). If fibrinogen were required, removal of fibrinogen from our perfusion system would have diminished the number of leukocyte interactions with primed platelets/α_IIb_β_3_ if Mac-1 were involved. 3) Antibody-mediated blocking of β_2_ integrins did not reduce VWF-’primed’ platelet-leukocyte interactions (**Fig 4c**). 4) Only neutrophils and T cells interact with the ‘primed’ platelets, whereas Mac-1 and LFA-1 are also highly expressed in monocytes which do not bind (**Fig 4d-f**).

We also excluded a role for P-selectin-PSGL-1 for the platelet-leukocyte interaction that we observe as; we detected little/no P-selectin on VWF-’primed’ platelets (**Fig S3**), suggestive of minimal degranulation occurring; this also provides indirect evidence for the lack of CD40L on the platelet surface. 2) P-selectin blockade had no effect upon the number of leukocytes binding (**Fig 3e**), and 3) only T-cells and neutrophils bind VWF-’primed’ platelets (**Fig 4**) - given that all leukocytes express PSGL-1,(Laszik et al., 1996) and CD40, if the capture of leukocytes were entirely P-selectin or CD40L-mediated, such cell-type selectivity would not be observed. We did however measure an influence of P-selectin upon the rolling speed of leukocytes over VWF-primed platelets (**Fig 3f & Fig S4d**). This suggests that although low levels of P-selectin present on the platelet surface is insufficient to facilitate leukocyte capture, it may synergize to slow rolling of leukocytes that are first captured by α_IIb_β_3_.

There are several studies that provide support for P-selectin-independent interactions of neutrophils and T-cells with platelets. Guidotti *et al* demonstrated the interaction of T-cells with small intrasinusoidal platelet aggregates in the liver during hepatotropic viral infections was independent of both P-selectin and CD40L in platelets (Guidotti et al., 2015). Using a murine model of peritonitis, Petri *et al* demonstrated that neutrophil recruitment and extravasation was highly dependent upon VWF, GPIbα, and platelets, but largely independent of P-selectin (Petri et al., 2010). Two further studies also corroborate the contention that VWF/GPIbα-bound platelets are capable of promoting neutrophil recruitment/extravasation in murine models of ischemia/reperfusion via P-selectin-independent mechanisms (Gandhi et al., 2012; Khan et al., 2012). These studies support the idea that both VWF and platelets can function beyond hemostasis to fulfil a role in leukocyte recruitment at sites of inflammation. As T-cells and neutrophils (and not B-cells or monocytes) can bind platelets via activated α_IIb_β_3_, this suggests that a specific receptor exists on these cells that is absent on B-cells or monocytes. Using transcriptomic and proteomic data, we identified 30 transmembrane candidates that were preferentially expressed in neutrophils (or T-cells) over monocytes. From this list, *SLC44A2* stood out due to its recent identification as a risk locus for both VTE and stroke, but with as yet unknown functional association with these pathologies (Apipongrat et al., 2019; Germain et al., 2015; Hinds et al., 2016). As platelet-leukocyte interactions are involved in both of these thrombotic disorders, we hypothesized that SLC44A2 functions as the neutrophil counter receptor for activated α_IIb_β_3_. The cellular function of SLC44A2 is not well-defined. It contains ten transmembrane domains with five extracellular loops. The intracellular N-terminal tail contains several putative phosphorylation sites of unknown functional significance. As well as neutrophils, SLC44A2 expression has also been reported in endothelial cells and platelets. However, proteomic data suggest that levels in neutrophils are >300 fold greater in neutrophils that platelets (Rieckmann et al., 2017).

We provide several lines of evidence to support the direct interaction between SLC44A2 and activated α_IIb_β_3_. 1) two different anti-SLC44A2 antibodies that recognize extracellular loops of the receptor blocked the binding of neutrophils to both VWF-’primed’ platelets and to activated α_IIb_β_3_. 2) recombinant expression of SLC44A2 in HEK293T cells imparted the ability of these cells to bind both VWF-primed platelets and activated α_IIb_β_3_ under flow in a manner that can be blocked by either GR144053 or by anti-SLC44A2 antibodies. 3) introduction of the rs2288904-A SNP in *SLC44A2* that is protective against VTE resulted in markedly reduced binding of transfected HEK293T cells to both VWF-’primed’ platelets and activated α_IIb_β_3_. 4) neutrophils homozygous for the rs2288904-A/A SNP exhibit significantly reduced binding to VWF-’primed’ platelets.

Although NET production is an established mechanism through which neutrophils control pathogens (Brinkmann et al., 2004), many questions remain as to how NETosis is regulated (Nauseef and Kubes, 2016). Binding of platelets to Kupffer cells in the liver of mice following infection with *B. cereus* or *S. aureus* is mediated by VWF (Wong et al., 2013). This binding augments the recruitment of neutrophils, NET production and the control of infection (Kolaczkowska et al., 2015). Mice lacking VWF or GPIbα do not form these aggregates and so have diminished neutrophil recruitment and, therefore, decreased survival (Wong et al., 2013). How NETosis is initiated following platelet binding remains uncertain. Alone, lipopolysaccharide (LPS) is not a potent activator of NETosis (Clark et al., 2007). However, LPS-stimulated platelets, which bind of fibrinogen (i.e. α_IIb_β_3_ is activated) and also robustly activate NETosis independent of P-selectin (Clark et al., 2007; Looney et al., 2009; Lopes Pires et al., 2017; McDonald et al., 2012).

We detected rapid release of intracellular Ca^2+^ (within minutes) in bound neutrophils (**Fig 5 & Movie 5**) that preceded the release of NETs after 80-90 minutes (**Fig 6**). This process was dependent upon neutrophils being captured under flow, suggesting that signal transduction through binding of α_IIb_β_3_ to SLC44A2 may be mechanosensitve, which is consistent with the recent report suggesting a major influence of shear upon NETosis in the presence of platelets (Yu et al., 2018). As NETosis can be induced following binding to purified α_IIb_β_3_ under flow alone, this suggests that this process does not require a component of the platelet releasate (e.g. mobility group box 1, platelet factor 4, RANTES and thromboxane A2) which have been reported to be capable of driving NETosis (Carestia et al., 2016). We propose a model in which platelets have a tunable response that can distinguish their roles in hemostasis and immune cell activation (**Fig 9**). The ‘priming’ of platelets by binding to VWF under flow (in the absence of other platelet agonists) may assist in the targeting of leukocytes to resolve pathogens or mediate vascular inflammatory response. The activated α_IIb_β_3_ integrin can then mediate neutrophil recruitment through binding to SLC44A2 (**Fig 9**). We do not exclude a supporting role for P-selectin in maintaining T-cell/neutrophil recruitment, but this is not essential for initiating recruitment. Under flow SLC44A2 transduces a mechanical stimulus capable of promoting NETosis via a pathway involving synergy between NADPH oxidase and Ca^2+^ signaling (**Fig 9**). Homeostatically, this may be beneficial for immune responses. However, during chronic infection or vascular inflammation NET production may promote intravascular thrombosis.

**Figure 9:**
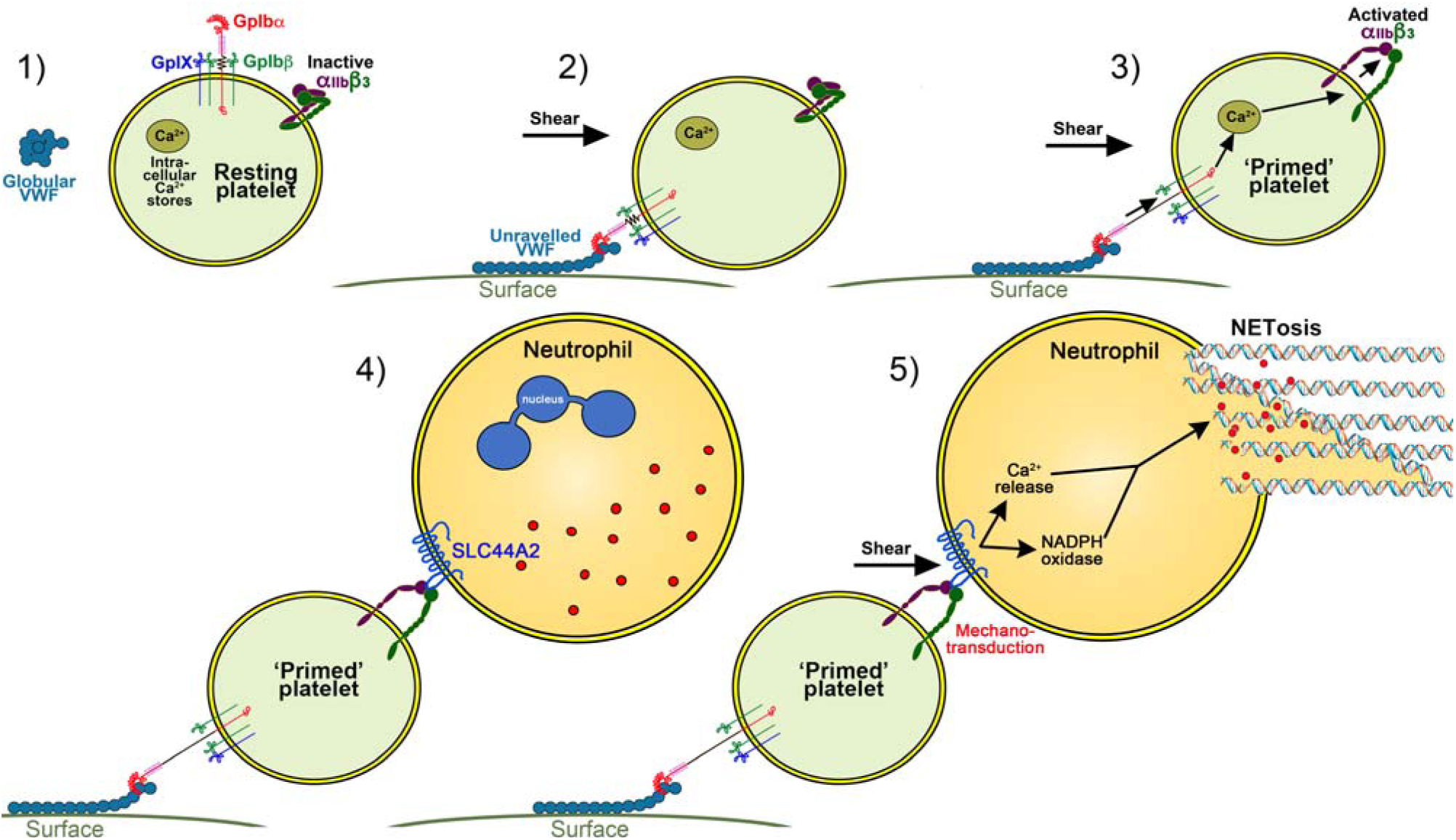
Model of platelet priming, neutrophil binding and NETosis. **1)** Under normal conditions, VWF circulates in plasma in a globular conformation that does not interact with platelets. Resting platelets present GPIbα on their surface - in complex with GPIbβ, GPIX and GPV - and also α_IIb_β_3_ in its inactive conformation. **2)** When VWF is attached to a cell surface (e.g. activated endothelial cell/Kuppfer cell or to a bacterial cell) or to an exposed collagen surface under flow, VWF unravels to expose its A1 domain enabling capture of platelets via GPIbα. **3)** Binding of platelets to VWF under flow induces mechano-unfolding of the juxtamembrane stalk of GPIbα leading to intraplatelet signaling, release of intraplatelet Ca^2+^ stores and activation of integrin α_IIb_β_3._ **4)** Neutrophils can bind to activated α_IIb_β_3_ under flow via SLC44A2. **5)** Shear forces on the neutrophil induce mechanosensitive signaling into the neutrophil causing intracellular Ca^2+^ release and NADPH oxidase-dependent NETosis.

This study identifies activated α_IIb_β_3_ as a receptor and agonist for neutrophils through SLC44A2. This provides a previously uncharacterized mechanism of how platelet-neutrophil cross-talk is manifest in innate immunity; it also provides an explanation for how VWF and platelet-dependent neutrophil recruitment and NETosis may occur in thrombotic disorders such as DVT (Laridan et al., 2019), but also thrombotic microangiopathies like thrombotic thrombocytopenic purpura (Fuchs et al., 2012b). Identification of SLC44A2 as the counter-receptor for activated α_IIb_β_3_ in conjunction with the prior identification of a protective SNP in SLC44A2 that impairs the binding of neutrophils to platelets highlights SLC44A2 as a potential therapeutic target. Recent data reveal that SLC44A2-deficient mice exhibit normal hemostatic responses (Tilburg et al., 2018), but are protected against development of venous thrombosis (Maracle et al., 2019), provide further encouragement for this strategy.

## Materials and Methods

### Preparation of VWF A1 domain and multimeric VWF

The coding sequence for the human VWF A1 domain (Glu1264 to Leu1469) was cloned into the pMT-puro vector, containing a C-terminal V5 and polyhistidine tag. The Y1271C/C1272R mutations were introduced by site-directed mutagenesis into A1 domain (A1*) (Blenner et al., 2014). All vectors were verified by sequencing.

Stably expressing S2 insect cells were selected using puromycin (Life Technologies). Cells were cultured under sterile conditions at 28°C in Schneider’s Drosophila medium (Lonza), supplemented with 10% heat-inactivated fetal bovine serum (FBS), 50μg/ml penicillin and 50U/ml streptomycin. Cells were grown in suspension in 2L conical flasks to a density of 2×10^6^ cells/ml. Expression of VWF A1 or A1* was induced by addition of 500μM CuSO_4_ for 5-7 days, at 28°C and 110 rpm.

Conditioned media were harvested, cleared by centrifugation, concentrated by tangential flow filtration and dialyzed against 20mM Tris (pH 7.8) 500mM NaCl. VWF A1 or A1* were purified by a two-step purification method using a Ni^2+^-HiTrap column followed by a heparin-Sepharose column (GE Healthcare) and elution with 20mM Tris, 600mM NaCl. VWF A1 and A1* were dialyzed in phosphate-buffered saline (PBS). A1 and A1* concentrations were determined by absorbance at 280nm. Proteins were analyzed by SDS-PAGE under reducing and non-reducing, and by Western Blotting using anti-His or anti-VWF antibodies. Full length, multimeric VWF was isolated from Haemate P by gel filtration and quantified by a specific VWF ELISA, as previously described (O’Donnell et al., 2005).

### Blood collection and processing

Fresh blood was collected in 40µM PPACK (for whole blood experiments), 3.13% citrate (for leukocyte isolation) or 85mM sodium citrate, 65mM citric acid, 111mM D(+) glucose, pH 4.5 (1x ACD, for plasma-free blood preparation). For reconstituted plasma-free blood, red blood cells (RBCs) and leukocytes were pelleted and washed twice. Separately, platelets were washed twice in 1x HEPES-Tyrode (HT) buffer containing 0.35% BSA, 75mU apyrase and 100nM prostaglandin E1 (Sigma). RBCs, leukocytes and platelets were resuspended in 1x HT buffer supplemented with 0.35% BSA. In some experiments, 1.3mg/ml purified fibrinogen (Haem Tech) was added. For Ca^2+^ assays, PRP was incubated with 5µM Fluo-4 AM (Thermo Fisher Scientific) for 30 minutes at 37°C prior to washing, and plasma-free blood was recalcified with 1mM CaCl_2_ (final concentration) immediately prior to flow experiments.

Polymorphonuclear cells (PMNs) and peripheral blood mononuclear cells (PBMCs) separated using Histopaque1077 and Histopaque1119 were resuspended in 1x HT, supplemented with 1.5mM CaCl_2_. For Ca^2+^ assays, PMNs were preloaded with 1µM Fluo-4 AM for 30 minutes at 37°C, before washing. This study was approved by the Imperial College Research Ethics Committee, and informed consent was obtained from all healthy volunteers.

### Flow experiments

VenaFluoro8+ microchips (Cellix) were coated directly with 2μM VWF in PBS overnight at 4°C in a humidified chamber. Coated channels were blocked for 1 hour with 1x HEPES-Tyrode (HT) buffer containing 1% bovine serum albumin (BSA). For the isolated VWF A1 and A1* domains, NTA PEGylated microchips (Cellix) were used to capture the A1 or A1* via their His tags (Tischer et al., 2014). Channels were stripped with EDTA before application of Co^2+^ and washing with 20mM HEPES, 150mM NaCl, pH 7.4 (HBS). To each channel, 20μl of 3.75μM VWF A1 or A1* were applied at room temperature for 20 minutes in a humidified chamber. Channels were then incubated with H_2_O_2_ for 30 min to oxidize Co^2+^ to Co^3+^, which stabilizes the binding of His-tagged A1/A1* (Wegner et al., 2016).

To NHS-microchannels (Cellix), 2.6μM purified α_IIb_β_3_ (ERL), 0.25mg/ml PECAM-1 (BioLegend), anti-β_3_/LIBS2 antibody (Millipore), anti-CD16 (eBiosciences) or 0.25mg/ml BSA were covalently attached by amine-coupling according to manufacturer’s instructions. For directly-coated α_IIb_β_3_ channels, the surface was washed with HBS containing 1mM MnCl_2_, 0.1mM CaCl_2_ following coating. Mn^2+^ was maintained in all subsequent buffers to cause α_IIb_β_3_ to favor its open, ligand binding conformation, as previously reported (Litvinov et al., 2005).

To anti-β_3_/LIBS2 antibody coated channels, α_IIb_β_3_ (ERL) was perfused over the surface to facilitate both capture and activation of α_IIb_β_3_ on the surface.

Whole blood or plasma-free blood was perfused through channels coated with either FL-VWF, A1, A1* or anti-PECAM-1 at shear rates of 500-1500s^-1^ for 3.5 minutes, followed by 50s^-1^ for 15 minutes using a Mirus Evo Nanopump and Venaflux64 software (Cellix). In separate experiments, 2.4μM eptifibatide (Sigma), 2μM GR144053 (Tocris), or 50μg/ml anti-P-selectin blocking antibody (clone AK4; BD Biosciences) were supplemented to whole blood or plasma-free blood. DiOC_6_ (2.5μM; Invitrogen) was used to label platelets and leukocytes. Cells were monitored in real-time using an inverted fluorescent microscope (Zeiss) or a SP5 confocal microscope (Leica). Leukocytes and platelets were distinguished by their larger size. For presentation and counting purposes, leukocytes were pseudo-colored to distinguish them. In some experiments, antibodies that recognize the second extracellular loop of SLC44A2, rabbit anti-SLC44A2 #1 (Abcam; Ab177877) or the first extracellular loop, rabbit anti-SLC44A2 #2 (LS Bio; LS-C750149) (0-20μg ml^-1^) to block SLC44A2 were compared to non-immune rabbit IgG (Abcam; 20μg ml^-1^) to explore the influence of SLC44A2 on neutrophils to bind to either VWF-‘primed’ platelets or isolated/activated α_IIb_β_3_.

Isolated PMNs and PBMCs were perfused through channels coated either directly or indirectly with α_IIb_β_3_, or BSA at 50s^-1^ for 15 minutes. Antibodies specific to the different types of leukocytes were added to isolated leukocytes, i.e. anti-CD16 (eBiosciences) conjugated to allophycocyanin (APC) to identify neutrophils, anti-CD14-APC for monocytes, anti-CD3-APC for T-cells and anti-CD19-APC for B-cells (BioLegend).

To visualize NETosis, neutrophils were labelled with 8μM Hoechst dye (cell permeable) and 1μM Sytox Green (cell impermeable) and monitored for 2 hours. As indicated, isolated PMNs were preincubated with 20μM TMB-8 (Ca^2+^ antagonist and protein kinase C inhibitor; Sigma), for 15 minutes, or 30μM DPI (NADPH oxidase inhibitor; Sigma) for 30 minutes at 37°C prior to NETosis assays. In some experiments, neutrophils were captured on microchannels coated with anti-CD16 and stimulated with 160nM PMA prior to analysis of NETosis in the presence and absence of inhibitors.

To confirm the presence of NETs, neutrophils that were captured by activated α_IIb_β_3_ and fixed with 4% paraformaldehyde after 2 hours. Fixed neutrophils were permeabilized with 0.1% Triton X-100 in PBS for 10 minutes, blocked with 3% BSA in PBS and, thereafter, incubated with rabbit polyclonal anti-citrullinated H3 (Abcam, 10μg/ml) overnight at 4°C (Martinod et al., 2013). Neutrophils were incubated with a goat anti-rabbit secondary antibody conjugated with Alexa647 (Abcam, 1:500) and with the Hoechst dye (8μM) for 2 hours, washed and then visualised by confocal microscopy.

Quantitation of platelet rolling, aggregation and intracellular Ca^2+^ release was achieved using SlideBook software (3i). The number of leukocytes rolling/attaching per minute at 50s^-1^ was derived by counting the number of cells in one field of view over a period of 13 minutes. NETosis was quantified by determining the proportion of all neutrophils in the microchannel that had undergone NETosis after 2 hours.

### Transcriptomic profiling of human leukocytes

RNA sequencing data from different leukocytes were obtained from the BLUEPRINT consortium [https://www.biorxiv.org/content/10.1101/764613v1]. For this, neutrophils and monocytes were isolated from peripheral blood. PBMCs were separated by gradient centrifugation (Percoll 1.078 g/ml) whilst neutrophils were isolated by CD16 positive selection (Miltenyi) from the pellet, after red blood cell lysis. PBMCs were further separated to obtain a monocyte rich layer using a second gradient (Percoll 1.066 g/ml) and monocytes further purified by CD14 positive selection (Miltenyi) after CD16 depletion. For neutrophils and monocytes, gene expression was tested also on Illumina HT12v4 arrays (accession E-MTAB-1573 at arrayexpress). The purification of naive B lymphocytes, naive CD4 lymphocytes, naive CD8 lymphocytes used in this study has been extensively described. Regulatory CD4 lymphocytes (T regs) were isolated by flow activated cytometry using the following surface markers combinations: CD3+ CD4+ CD25+ CD127low. Cell type purity was assessed by flow cytometry and morphological analysis. RNA was extracted using TRIzol according to manufacturer’s instructions, quantified using a Qubit RNA HS kit (Thermofisher) and quality controlled using a Bioanalyzer RNA pico kit (Agilent). For all cell types libraries were prepared using a TruSeq Stranded Total RNA Kit with Ribo-Zero Gold (Illumina) using 200ng of RNA. Trim Galore (v0.3.7) (http://www.bioinformatics.babraham.ac.uk/projects/trim_galore/) with parameters “-q 15 -s 3 --length 30 -e 0.05” was used to trim PCR and sequencing adapters. Trimmed reads were aligned to the Ensembl v75 human transcriptome with Bowtie 1.0.1 using the parameters “-a --best --strata -S -m 100 -X 500 --chunkmbs 256 --nofw --fr”. MMSEQ (v1.0.10) was used with default parameters to quantify and normalize gene expression. Differential gene expression analyses were performed: mature neutrophils (n=7) vs monocytes (n=5) and CD4-positive/αβ T cells (n=8) vs monocytes (n=5). Regulatory T cells (T_reg_, n=1) and native B cells (n=1), are included in the heatmap, for comparison but were not used in differential gene expression analysis due to the low number of biological replicates. We selected genes that were expressed significantly higher in neutrophils than in monocytes, and also those that were significantly higher in CD4-positive/αβ T cells than in monocytes. Their intersection identified 750 genes (598 of which protein coding). From these 598 genes, we selected the 93 genes that contained the Uniprot annotation of “INTRAMEMBRANE DOMAIN” or “TRANSMEM DOMAIN”. The effective log2(FPKM+1) data were presented in the heatmap. Further selection involved discarding those transmembrane proteins that are not present on the extracellular membrane, or primarily associated with intracellular membranes as determined by Uniprot annotation. Proteins that (where known) had extracellular regions of <30 amino acids, as determined in Uniprot, that might be less likely capable of mediating specific ligand binding were also excluded. Finally, analysis of proteomic data from the ImmProt (http://immprot.org) resource was used to verify higher levels of protein of each selected gene in neutrophils than in monocytes

### Expression of SLC44A2 in HEK293T cells

The mammalian expression vector, pCMV6-Entry containing the human *SLC44A2* cDNA C-terminally fused to EGFP was purchased from OriGene. To introduce the rs2288904 SNP encoding a R154Q substitution, site-directed mutagenesis was performed using the primers; 5’-GTG GCT GAG GTG CTT CAA GAT GGT GAC TGC CCT-3’, and 5’-AGG GCA GTC ACC ATC TTG AAG CAC CTC AGC CAC-3’. Successful introduction of the SNP was confirmed by sequencing.

HEK293T cells were cultured as adherent layers, in humidified incubators at 37°C, 5% CO_2_, in minimum essential media (MEM; Sigma) supplemented with 10% FBS, 1U/ml Penicillin 0.1mg/ml Streptomycin, 1% non-essential amino acids (Sigma) and 2mM L-glutamine.

Cells were seeded in 6-well plates 24 hours prior to transfection and transfected using Lipofectamine 2000 (Invitrogen). Transfection efficiency was visually observed using fluorescent microscopy and quantified using flow cytometry. In all cases transfection efficiency was >75%. Cells were harvested 24 hours post-transfection with Tryplex (Life Tech) to obtain a single cell suspension. Cells were washed with complete medium and cells resuspended in serum-free OptiMEM (Life Tech) until use.

### Flow assays using HEK293T cells

Microchannels were coated with FL-VWF or α_IIb_β_3_ (coupled via the anti-β_3_/LIBS2 antibody). Thereafter, unlabeled plasma-free blood was perfused over the FL-VWF coated channels at high shear for 3.5 minutes to capture a layer of ‘primed’ platelets. Platelet coverage was monitored in bright-field. Channels were subsequently washed with 1xHT buffer to remove the blood and SLC44A2-EGFP transfected HEK293T cells were perfused at low shear (25s^-1^) for 10 minutes. Transfected HEK293T cells were also perfused through FL-VWF coated channels (in the absence of platelets for 30 min at 25s^-1^) to examine any direct interaction with VWF. Transfected HEK293T cells were also perfused through α_IIb_β_3_ (coupled via the anti-β_3_/LIBS2 antibody) channels at 25s^-1^ for 10 minutes. Binding of EGFP fluorescent HEK293T cells was quantified by counting the number of cells attached after 10 minutes across the whole channel and then expressing this as the mean number of cells/field of view. In separate experiments, the ability of GR144053 to block α_IIb_β_3_, or antibodies that recognize the first extracellular loop of SLC44A2, rabbit anti-SLC44A2 #1 (Abcam; Ab177877) or rabbit anti-SLC44A2 #2 (LS Bio; LS-C750149) (0-20μg ml^-1^) to block SLC44A2 were compared to non-immune rabbit IgG (Abcam; 20μg ml^-1^)

### Genotyping

To identify individuals homozygous for the *SLC44A2* rs2288904-A SNP or for the wild type allele rs2288904-G, 25μl blood was taken by pin prick from healthy volunteers that provided written informed consent. Genomic DNA was then extracted using PureLink® Genomic DNA kit (Invitrogen). DNA yield was quantified by NanoDrop. Genomic DNA from each volunteer was used as a template to PCR amplify a 410 base pair fragment of the *SLC44A2* gene spanning the SNP site using primers 5’-ACC TCA CGT ACC TGA ATG-3’ and 5’-AGC CAT GCC CAT CCT CAT AG-3’. After amplification, samples were separated by agarose gel electrophoresis and the 410 bands excised, purified using the Gel Extraction kit (Qiagen) and sequenced using the first PCR primer. PMN isolated from genotyped individuals were subsequently used to examine their ability to bind both activated α_IIb_β_3_ (captured using the LIBS2, anti-β_3_ antibody) and VWF-’primed’ platelets, as described above.

### Statistics

Statistical analysis was performed using Prism 6.0 software (GraphPad). Differences between data/samples was analyzed using unpaired two-tailed Student’s t-test or Mann-Whitney, as appropriate and as indicated in figure legends. Data are presented as mean ±standard deviation, or median ± 95% confidence interval. The number of individual experiments performed (n) is given in each legend. Values of p<0.05 were considered statistically significant.

### Data availability

Data supporting the findings of the study are available from the corresponding author upon reasonable request. The source data underlying Figs 1c, 2b-e, 3, 4b-c, 5c-d, 6c-e, 8a, c, d and f and Supplementary Figs 1, 2, 3c-d, 4b are available upon request.

## Supporting information

Suuplemntary data

## Acknowledgements

This work was funded through by grants from the British Heart Foundation (FS/15/65/32036 and PG/17/22/32868) awarded to J.T.B.C, K.J.W and I.I.S-C. M.F is supported by the British Heart Foundation (FS/18/53/33863). The authors declare no competing financial interests. The authors would like to thank Ying Jin and My Dang (imperial College London) for technical assistance and blood sampling. The authors would also like to thank Prof Heyu Ni and Dr Miguel Neves (University of Toronto) for helpful discussions into identification of α_IIb_β_3_ binding partners.

## Author contributions

A.C-B designed and performed the experiments, analyzed the data, prepared the figures and wrote the manuscript; I.I.S-C designed and performed the experiments, analyzed the data, and wrote the manuscript L.G and M.F performed leukocyte transcriptional profiling experiments. K.J.W designed experiments, analyzed the data, and wrote the manuscript; J.T.B.C designed experiments, analyzed the data, prepared the figures and wrote the manuscript.

