## Supplementary material for "Activated α_IIb_β_3_ on platelets mediates flow-dependent NETosis via SLC44A2": Suuplemntary data

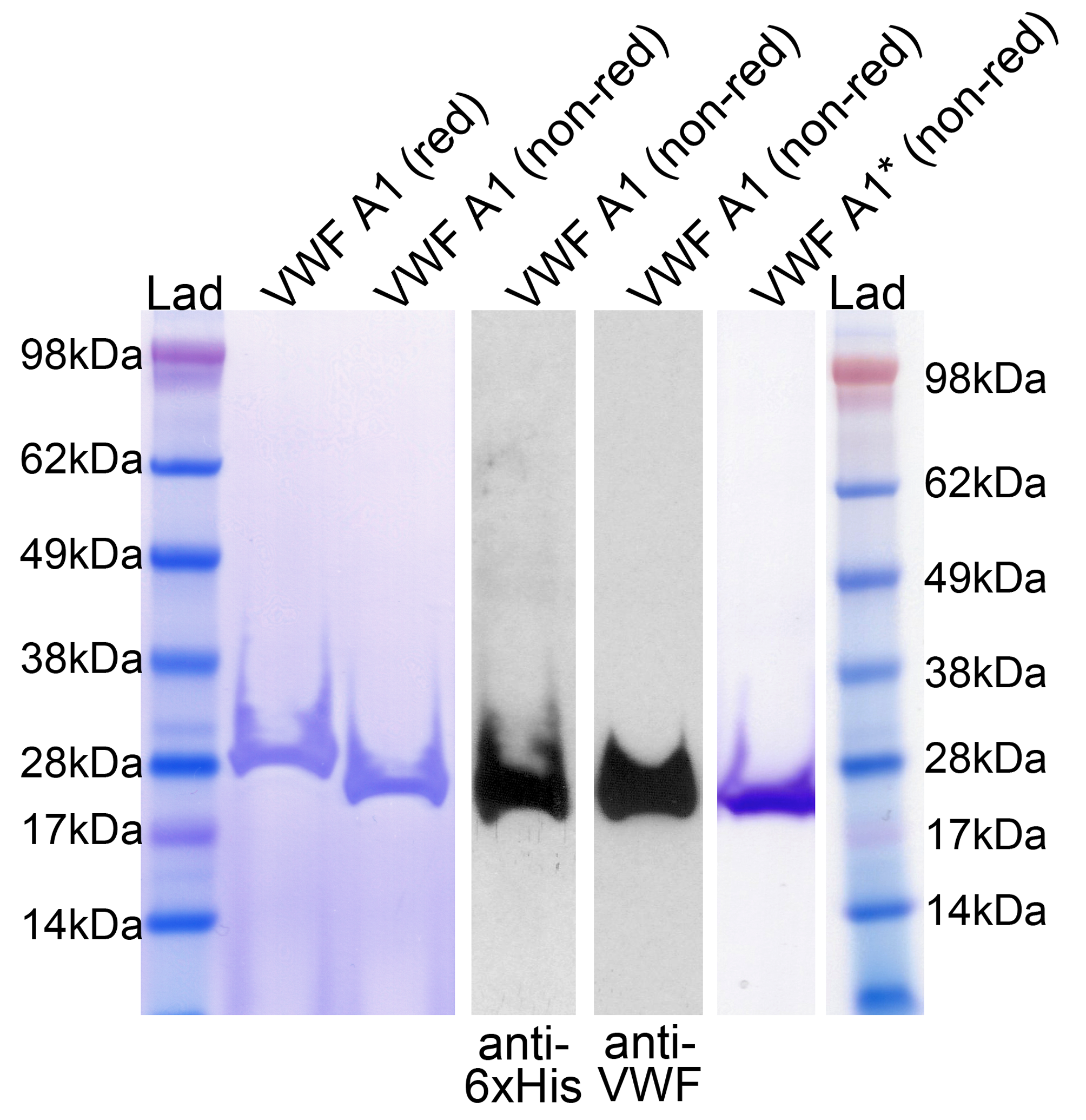

**Figure S1: Analysis of purified recombinant VWF A1 and VWF A1*.** VWF A1 domain with a C-terminal V5 and 6xHis tag was expressed in S2 insect cells and purified to homogeneity. Purified A1 was analyzed by SDS-PAGE and Coomassie staining under reducing and non-reducing conditions, and also by Western blotting of VWF A1 using anti-6xHis and anti-VWF antibodies. A Coomassie stain of purified VWF A1* is also shown.

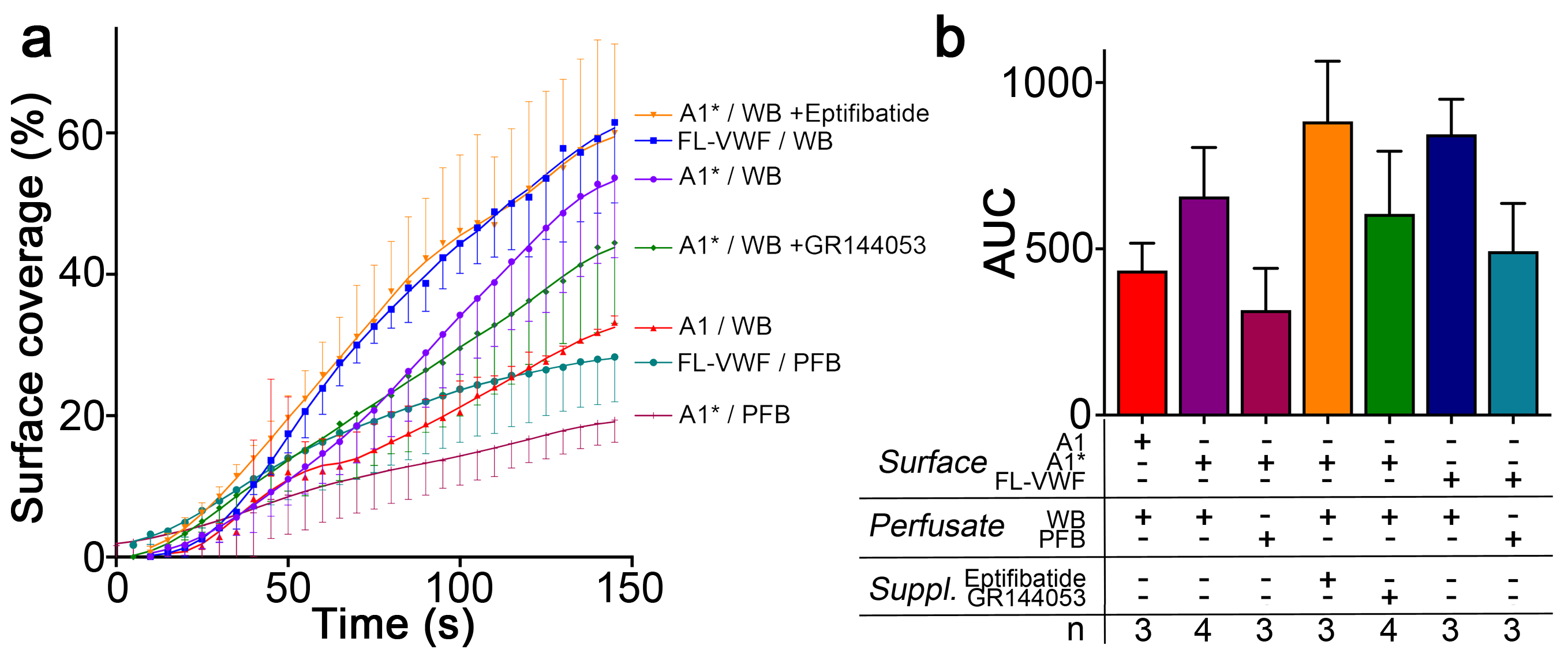

**Figure S2: Platelet coverage: a)** Graph showing the surface coverage over time of microchannels coated with FL-VWF, VWF A1 or VWF A1* by platelets in either whole blood (WB) or plasma-free blood (PFB) at a shear rate of 1000s^-1^ **b)** Graph of the area under the curve (AUC) ±SD of conditions analyzed in A) Surface coverage rates were similar on all surfaces.

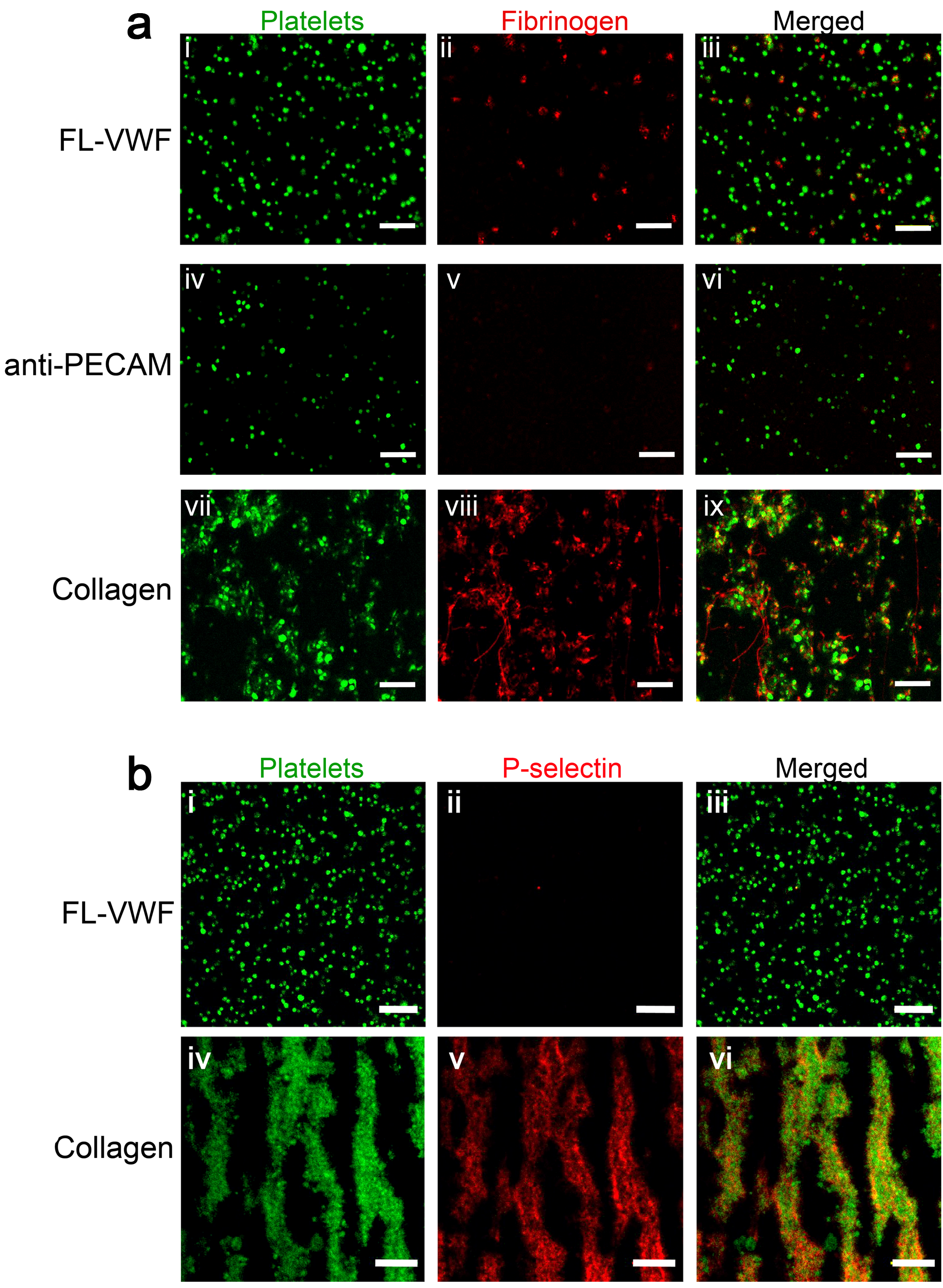

**Figure S3: Platelets binding to VWF under flow are ‘primed’ leading to activation of α_IIb_β_3_, but minimal presentation of P-selectin. a)** Representative images (n=3) depicting platelets (DiOC_6_, green) captured from plasma-free blood at 1000s^-1^ for 3.5 minutes onto microchannel surfaces coated with either FL-VWF (i-iii), anti-PECAM (iv-vi) or collagen (vii-ix). Plasma-free blood was supplemented with fibrinogen-Alexa647 (red). Merged images show fluorescent fibrinogen attached to the platelets captured by FL-VWF and collagen, but not by platelets captured by anti-PECAM. **b)** Representative images (n=3) depicting platelets (DiOC_6_, green) captured from whole blood at 1000s^-1^ for 3.5 minutes onto microchannel surfaces coated with either FL-VWF (i-iii) or collagen (iv-vi). Blood was supplemented with anti-P-selectin-APC (red). Merged images show the presence of P-selectin on the surface of platelets captured by collagen, but very little/no P-selectin on platelets captured FL-VWF. Scale bar=20μm.

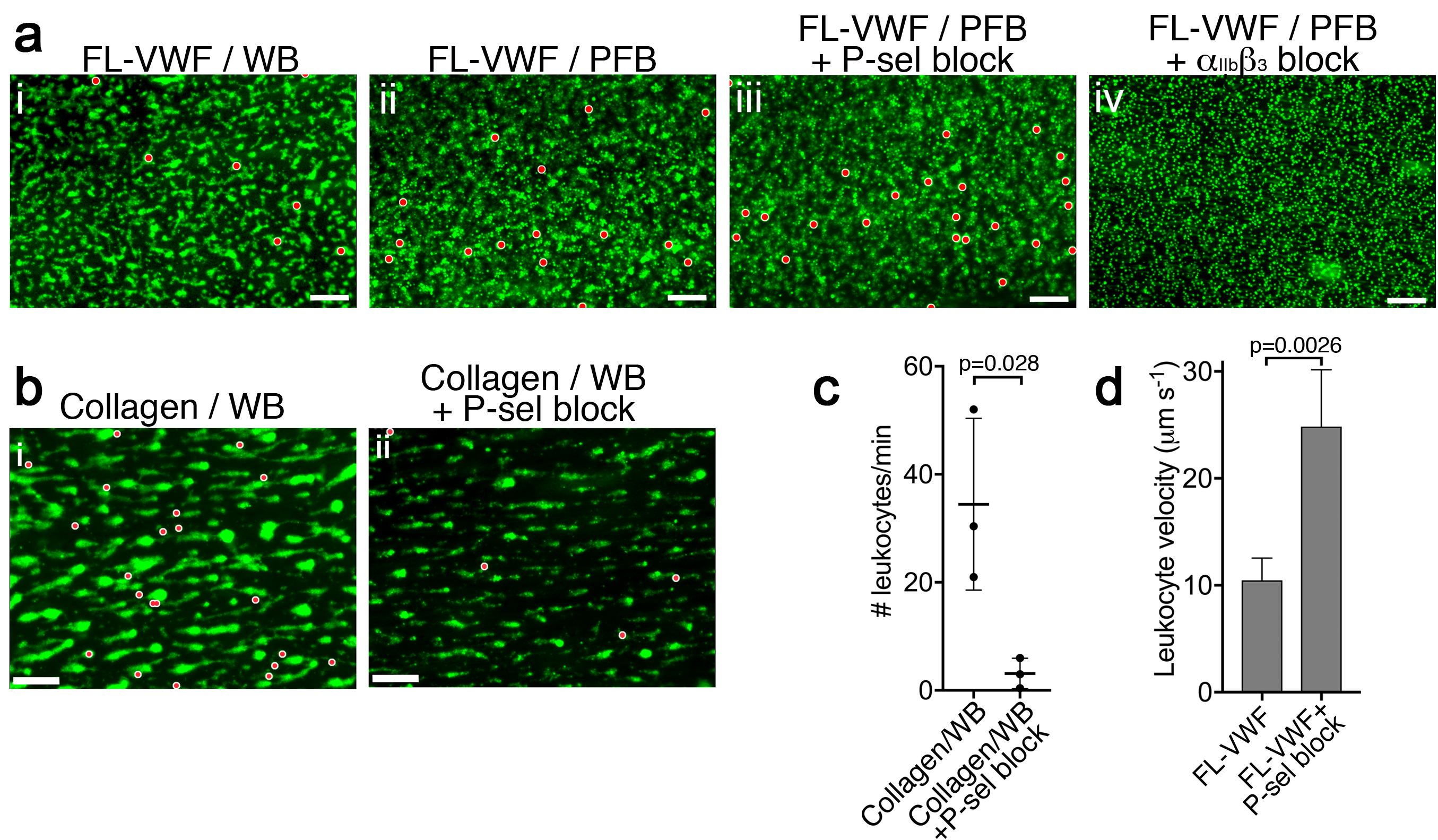

**Figure S4: Antibody-mediated blockade of P-selectin diminishes leukocyte binding to collagen bound platelets. a)** Vena8 microchannels were coated with FL-VWF. i) Whole blood (WB), ii) plasma-free blood (PFB), iii) WB containing anti-P-selectin blocking antibody, or iv) WB containing eptifibatide were perfused through channels at 1000s^-1^ for 3.5 minutes. Thereafter, the shear rate was reduced to 50s^-1^ for 5 minutes. Platelets and leukocytes were labelled with DiOC_6_, platelets are shown in green, leukocytes have been pseudo-colored red to aid visualization. Scale bar; 50μm (see also **Movie EV 2**). **b)** Whole blood (WB), labelled with DiOC_6_ was perfused over collagen microchannels at 1000s^-1^ for 3.5 minutes. Thereafter, the shear rate was reduced to 50s^-1^ to monitor leukocyte interactions. (i) Representative image (n=3) shown after 5 minutes platelets (green) and leukocytes (pseudo-colored red). (ii) as in (i) except WB was supplemented with a blocking anti-P-selectin antibody (AK-4 clone. Scale bar=50μm. **c)** The mean number of leukocytes binding per minute ±SD was calculated for each channel. There was a significant reduction in the number of leukocytes binding to collagen-bound platelets in the presence of P-selectin blocker, although leukocyte binding was not completely inhibited, suggesting that not all binding to collagen-activated platelets is P-selectin-dependent. **d)** Graph of leukocyte rolling velocity on platelets bound to FL-VWF in WB in the absence or presence of a blocking anti-P-selectin antibody. Data shown are the means of individual leukocyte rolling velocities for 3 separate experiments. Unpaired, two-tailed Student’s t test.

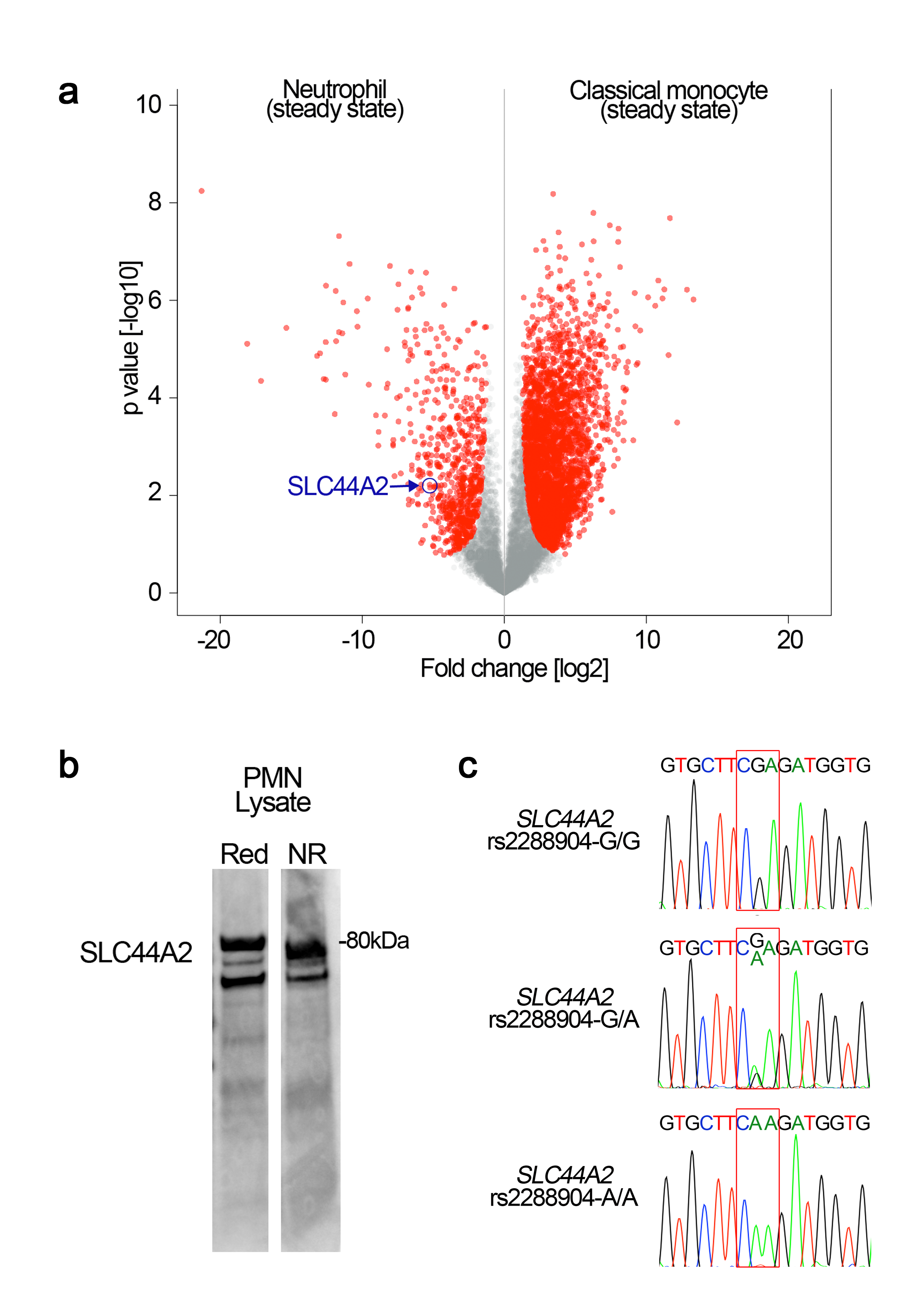

**Figure S5: SLC44A2 expression in neutrophils and *SLC44A2* genotype analysis.** **a)** Preferential expression of SLC44A2 in neutrophils over monocytes was ascertained through analysis of proteomic data from ImmProt (http://immprot.org), which is based on high-resolution mass-spectrometry proteomics performed on human hematopoietic cell populations in either steady or activated states. A pair-wise comparison between neutrophils (steady-state) and classical monocytes (steady state) is displayed as a volcano plot. Protein expression levels were compared using two-tailed Student’s t-test with Welch’s correction (S0=1, FDR<5%), and proteins with a significantly different expression level are shown in red on the plot. The point corresponding to SLC4A2 is highlighted revealing preferential expression in neutrophils. All 30 candidate genes identified in Fig 8 were analyzed through this comparative approach to examine preferential protein levels in neutorphils above monocytes. **b)** Isolated neutrophil lysates were analyzed by Western blotting under reducing and non-reducing conditions using anti-SLC4A2 #2 antibody. In both lanes, two bands corresponding to glycosylated (~80kDa) and nascent, non-glycosylated SLC44A2 are detected. **c)** Healthy volunteers were genotyped to identify individuals homozygous for the protective rs2288904-A/A SNP in *SLC44A2*. Representative chromatograms for individuals with rs2288904-G/G, rs2288904-G/A, rs2288904-A/A genotypes are shown.

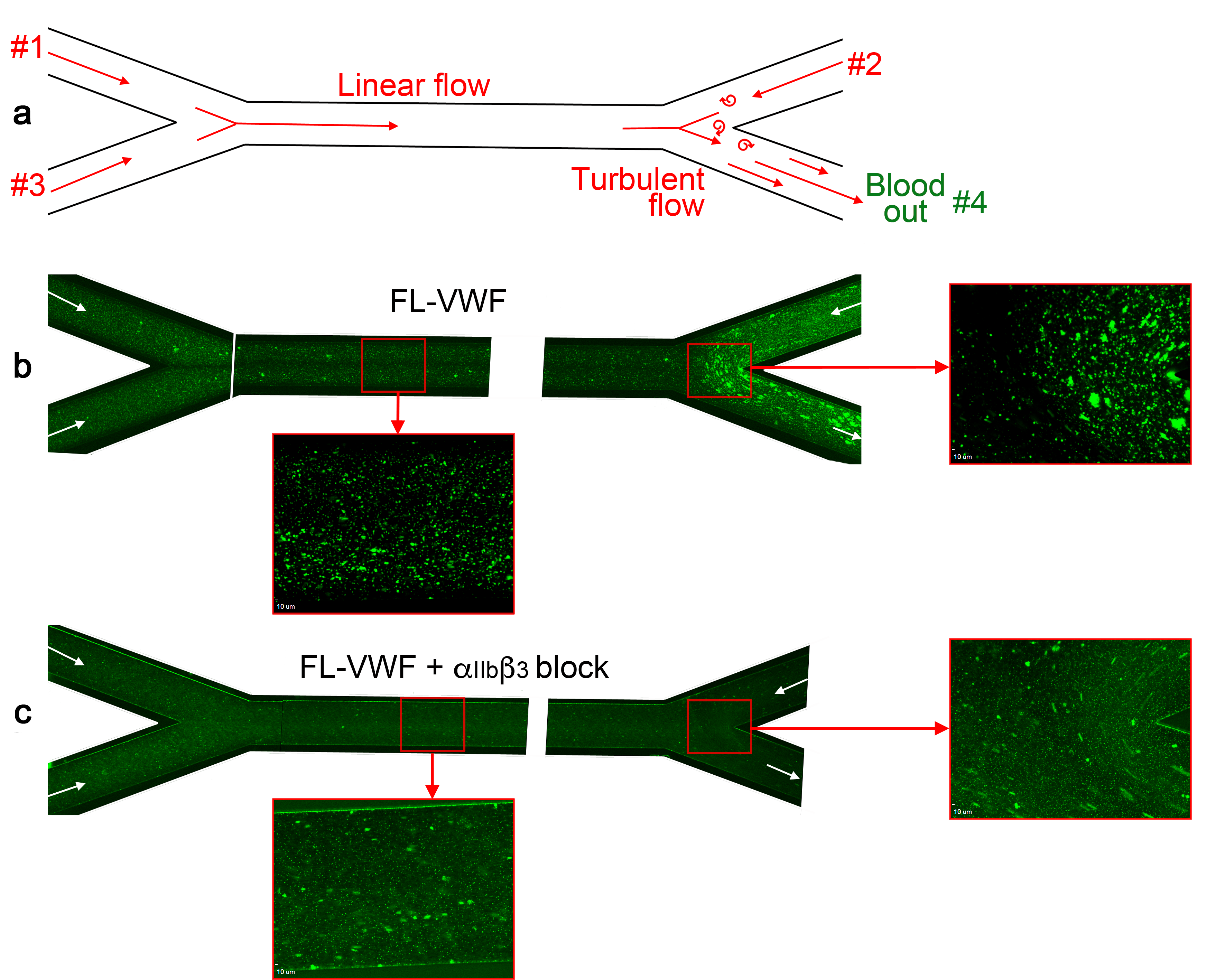

**Figure S6: Analysis of platelet binding to FL-VWF under low/disturbed flow and subsequent leukocyte binding. a)** Schematic representation of blood flow through bifurcated channels. Blood is drawn through inlets #1, #2 and #3, and out through outlet #4. For much of the channels, the flow is linear, with particular exception to the bifurcation site on the right where disturbed flow exists due to convergence of flows. **b)** Channels were coated with FL-VWF and whole blood labelled with DiOC6 was perfused through channels as in a), and as denoted by arrows, at an exit shear rate of 50s^-1^. At this low shear rate, platelets can bind to the channel surface to which leukocytes (larger cells also stained in green) also bind. Although this is evident in the linear part of the channel (inset), at the site of most turbulent flow increased platelet and leukocyte binding was observed. **c)** As in b) except GR144053 was added to block α_IIb_β_3_. Blocking α_IIb_β_3_ inhibited the majority of leukocyte binding to platelets (the majority of those observed are in transit). However, in the absence of leukocyte binding the binding of platelets to the VWF surface under low flow can be more clearly observed in both the linear and turbulent flow areas.

**Legends to Movies (see Movie 1-7 files)**

**Movie 1. Platelet capture on FL-VWF, VWF A1 or VWF A1*.** Vena8 microchannels were coated with either full-length VWF (FL-VWF), VWF A1 or A1*. Whole blood labelled with DiOC_6_ was perfused at 1000s^-1^ for 3 minutes. Note the reduced rate of platelet rolling on microchannels coated with A1* when compared to FL VWF or A1.

**Movie 2. VWF A1-GpIbα interaction induces intraplatelet Ca^2+^ release under flow.** Platelets were pre-loaded with Fluo-4 AM prior to perfusion through VWF A1*-coated microchannels at 1000s^-1^ for 5 minutes. Increases in platelet fluorescence corresponds to platelet intracellular Ca^2+^ release following attachment to VWF A1*. Note the repeated transient increases in fluorescence under flow, indicative of sustained and repeated signaling stimuli.

**Movie 3. Leukocytes bind/roll on VWF-‘primed’ platelets.** Whole blood or plasma-free blood labelled with DiOC_6_ was perfused through channels coated with VWF A1* at high shear for 3.5 minutes to capture platelets in the presence or absence of either a blocking anti-P-selectin antibody or eptifibatide (α_IIb_β_3_ blocker) prior to the acquisition of videos. The shear rate was reduced to 50s^-1^ to visualize leukocyte rolling or attaching (tracked in blue). Leukocytes rolled/bound on platelets in whole blood, plasma-free blood and plasma-free blood containing anti-P-selectin blocking antibody, but not in the presence of eptifibatide (α_IIb_β_3_ blocker). The use of the anti-P-selectin blocking antibody did however increase the rolling velocity of leukocytes over the platelet surface.

**Movie 4. Neutrophil start scanning following interaction with activated α_IIb_β_3_.** Following attachment of neutrophils (labelled with anti-CD16; red) to platelets (green) captured on FL-VWF (left), neutrophils start to scan the platelet surface. Similarly, neutrophils (anti-CD16-APC & Hoechst staining) bound to activated α_IIb_β_3_-coated microchannels also scanned the surface and in some cases appeared to start migrating against the direction of flow. Shear rate = 50s^-1^.

**Movie 5. Intracellular Ca^2+^ release in neutrophils following interaction with activated α_IIb_β_3_ under flow.** Neutrophils were pre-loaded with Fluo-4 AM and perfused over FL-VWF ‘primed’ platelets, α_IIb_β_3_-coated or anti-CD16 coated microchannels at 50s^-1^. Following attachment of neutrophils to VWF-‘primed’ platelets or activated α_IIb_β_3_ (but not anti-CD16), an increase in fluorescence corresponding to neutrophil intracellular Ca^2+^ release was observed.

**Movie 6. α_IIb_β_3_-induced NETosis.** Isolated PMNs labelled with Hoechst (blue) and cell-impermeable Sytox Green were perfused over α_IIb_β_3-_coated microchannels at 50s^-1^ for 10 minutes and then monitored under static conditions. Movie shown represents period from 80 to 95 minutes after attachment. Sytox Green staining appears, indicative of DNA becoming extracellular highlights the neutrophils undergoing NETosis.

**Movie 7. Neutrophil binding to VWF-‘primed’ platelets occurs via SLC44A2 is modified by the rs2288904 SNP.** Plasma-free blood generated from individuals homozygous for the rs2288904-G major allele in *SLC44A2* (SLC44A2 (R154/R154)) or the rs2288904-A minor allele (SLC44A2 (Q154/Q154)) SNP in *SLC44A2* was labelled with DiOC_6_ and perfused through channels coated with VWF at high shear for 3.5 minutes. Shear was subsequently reduced to 50s^-1^. SLC44A2 (R154/R154) leukocytes (tracked in blue) were seen to roll on the VWF-‘primed’ platelets. The number of leukocytes interacting was severely reduced in the presence of the anti-SLC44A2 #2 antibody. Leukocytes homozygous for the rs2288904-A minor allele, SLC44A2 (Q154/Q154), associated with protection against venous thrombosis, exhibited reduced ability to interact with VWF-‘primed’ platelets.
